# The ChAHP chromatin remodelling complex regulates neurodevelopmental disorder risk genes to scale the production of neocortical layers

**DOI:** 10.1101/2024.02.12.579820

**Authors:** Samuel Clémot-Dupont, José Alex Lourenço Fernandes, Sarah Larrigan, Xiaoqi Sun, Suma Medisetti, Rory Stanley, Ziyad El Hankouri, Shrilaxmi V. Joshi, David J. Picketts, Karthik Shekhar, Pierre Mattar

## Abstract

Although chromatin remodellers are among the most important risk genes associated with neurodevelopmental disorders (NDDs), the roles of these complexes during brain development are in many cases unclear. Here, we focused on the recently discovered ChAHP chromatin remodelling complex. The zinc finger and homeodomain transcription factor ADNP is a core subunit of this complex, and *de novo ADNP* mutations lead to intellectual disability and autism spectrum disorder. However, germline *Adnp* knockout mice were previously shown to exhibit early embryonic lethality, obscuring subsequent roles for the ChAHP complex in neurogenesis. Here, we employed single cell transcriptomics, cut&run-seq, and histological approaches to characterize mice conditionally ablated for the ChAHP subunits *Adnp* and *Chd4*. We show that during neocortical development, Adnp and Chd4 orchestrate the production of late-born, upper-layer neurons through a two-step process. First, Adnp is required to sustain progenitor proliferation specifically during the developmental window for upper-layer cortical neurogenesis. Accordingly, we found that Adnp recruits Chd4 to genes associated with progenitor proliferation. Second, in postmitotic differentiated neurons, we define a network of risk genes linked to NDDs that are regulated by Adnp and Chd4. Taken together, these data demonstrate that ChAHP is critical for driving the expansion upper-layer cortical neurons, and for regulating neuronal gene expression programs, suggesting that these processes may potentially contribute to NDD etiology.

**Highlights:**

- *Adnp* and *Chd4* cKOs exhibit similar deficits in cortical growth
- Adnp sustains the proliferation of apical progenitors to scale the production of upper-layer neurons
- Adnp recruits Chd4 to genes involved in corticogenesis
- Adnp is a master regulator of risk genes associated with neurodevelopmental disorders

## Introduction

During neurogenesis, developmental transitions in cell identity are accompanied by dynamic changes in gene expression. Chromatin remodelling complexes play a key role in facilitating such transitions by sliding or evicting nucleosomes to reconfigure gene regulatory elements. Possibly for this reason, chromatin remodellers are among the most commonly mutated genes associated with neurodevelopmental disorders (NDDs)^1^. However, in many cases, the underlying molecular mechanisms are poorly understood. The recently described ChAHP chromatin remodelling complex is a case in point. The core subunits of ChAHP are the ATP-dependent chromatin remodelling enzyme <u>CH</u>D4 (Mi-2β), the zinc finger and homeodomain transcription factor <u>A</u>DNP, and the <u>HP</u>1 (Cbx3 or Cbx5) heterochromatic proteins^2^. *ADNP* is among the most frequently *de novo* mutated genes associated with autism spectrum disorder (ASD), and is also linked to intellectual disability (ID)^3–8^. ADNP Syndrome - also known as Helsmoortel-Van der Aa syndrome, is typically driven by *de novo* heterozygous truncating mutations^9^. *CBX3/5* have not yet been linked to NDDs, but *de novo* mutations in *CHD4* are linked to Sifrim-Hitz-Weiss syndrome and ID^10–12^.

Despite the prominent genetic linkage between ChAHP subunits and NDDs, the precise role of the ChAHP complex in neurodevelopment has not yet been established. Indeed, ADNP, CHD4, and HP1 proteins have each been suggested to perform a variety of functions independently of ChAHP. For example, CHD4 is well known to be a key subunit in the <u>Nu</u>cleosome <u>R</u>emodelling and <u>D</u>eacetylase (NuRD) complex – particularly in neural progenitors^13^. HP1 proteins likewise interact with numerous other protein complexes in addition to ChAHP. ADNP has therefore been considered to be the only ChAHP-specific subunit, which is in agreement with data from unbiased proteomic studies^2,14,15^. However, ADNP has been shown to interact with additional protein partners – both in the nucleus^16–19^, and in the cytoplasm^20–25^. Such interactions could potentially allow ADNP to regulate neurodevelopment independently of ChAHP^3^.

Perhaps accordingly, phenotypes associated with *ADNP* and *CHD4* mutations are somewhat divergent. For example, in humans, macrocephaly is associated with *CHD4* mutations but not with *ADNP* mutations, and ASD is prominently linked to mutations in *ADNP* but not *CHD4*^6,9–12^. In mice, conditional knockouts (cKOs) have revealed a role for *Chd4* in several neurodevelopmental functions, including cortical growth^13,26–28^. By contrast, *Adnp* knockout mice die early in embryogenesis, and exhibit defects in neural progenitor induction^29^. In pluripotent cells, the competence to differentiate into neural progenitors was similarly lost when *ADNP* was mutated^2,20,30^. Importantly, the prominent genetic association with ASD/ID hints that ADNP likely regulates later neurodevelopmental processes, but appropriate model systems for deciphering such functions have thus far been lacking.

To examine how ChAHP regulates neurogenesis, we generated conditional knockout (cKO) mice in which *Adnp* or *Chd4* were equivalently ablated in the developing dorsal telencephalon. We found that *Adnp* and *Chd4* cKOs exhibited nearly identical reductions in brain size. Despite the fact that both Adnp and Chd4 proteins were expressed ubiquitously, we reveal a specific requirement for these genes in the expansion of upper-layer neocortical neurons. Next, using multiplexed single cell RNA-seq (scRNA-seq), we identified an overlapping transcriptomic signature that we attribute to ChAHP regulatory functions. Accordingly, we found that Adnp was required to recruit Chd4 to the regulatory elements of genes involved in cortical growth. Overall, our study identifies novel *in vivo* roles for the ChAHP complex during CNS development, and suggests a model where Adnp and Chd4 co-regulate gene expression to balance progenitor self-renewal and cortical expansion.

## Results

### Conditional ablation of ChAHP subunits in the developing telencephalon

During cortical development, excitatory glutamatergic projections neurons are produced by multipotent progenitor cells within the dorsal telencephalon. At approximately embryonic day (E) E11.5, apical progenitors (radial glia) located within the ventricular zone (VZ) begin to undergo asymmetric divisions that self-renew and additionally generate neuronal daughter cells. Beginning at ∼E12.5, apical progenitors also produce basal (intermediate) progenitors, which form a secondary progenitor layer called the subventricular zone (SVZ). Basal progenitors tend to divide symmetrically to generate neurons^31–34^. As they differentiate, each neuronal subtype migrates into a discrete layer within the cortical plate (CP) of the neocortex. In particular, neocortical subtypes are produced in an ordered but overlapping sequence, with deep-layer neurons produced first (subplate → Layer VI → Layer V), and upper-layer neurons produced secondarily (Layer IV → Layer II/III neurons (fused in mouse)). Neocortical progenitors are programmed to produce specific numbers of each neuronal subtype, although the mechanisms that scale cortical layers are not well understood.

Despite its prominent genetic association with NDDs, the functions of the ChAHP complex during cortical development remain unclear. We first examined the spatiotemporal expression pattern of Adnp in murine neocortical neurogenesis. We found that *Adnp* transcript and protein was robustly and ubiquitously expressed throughout murine brain development in all cell types (Fig. 1A-D; Fig. S1). However, we noted that in the adult brain, Adnp protein was expressed at higher levels in upper-layer cortical neurons (Fig. 1C, D; Fig. S1) as previously reported^35^.

**Figure 1.**
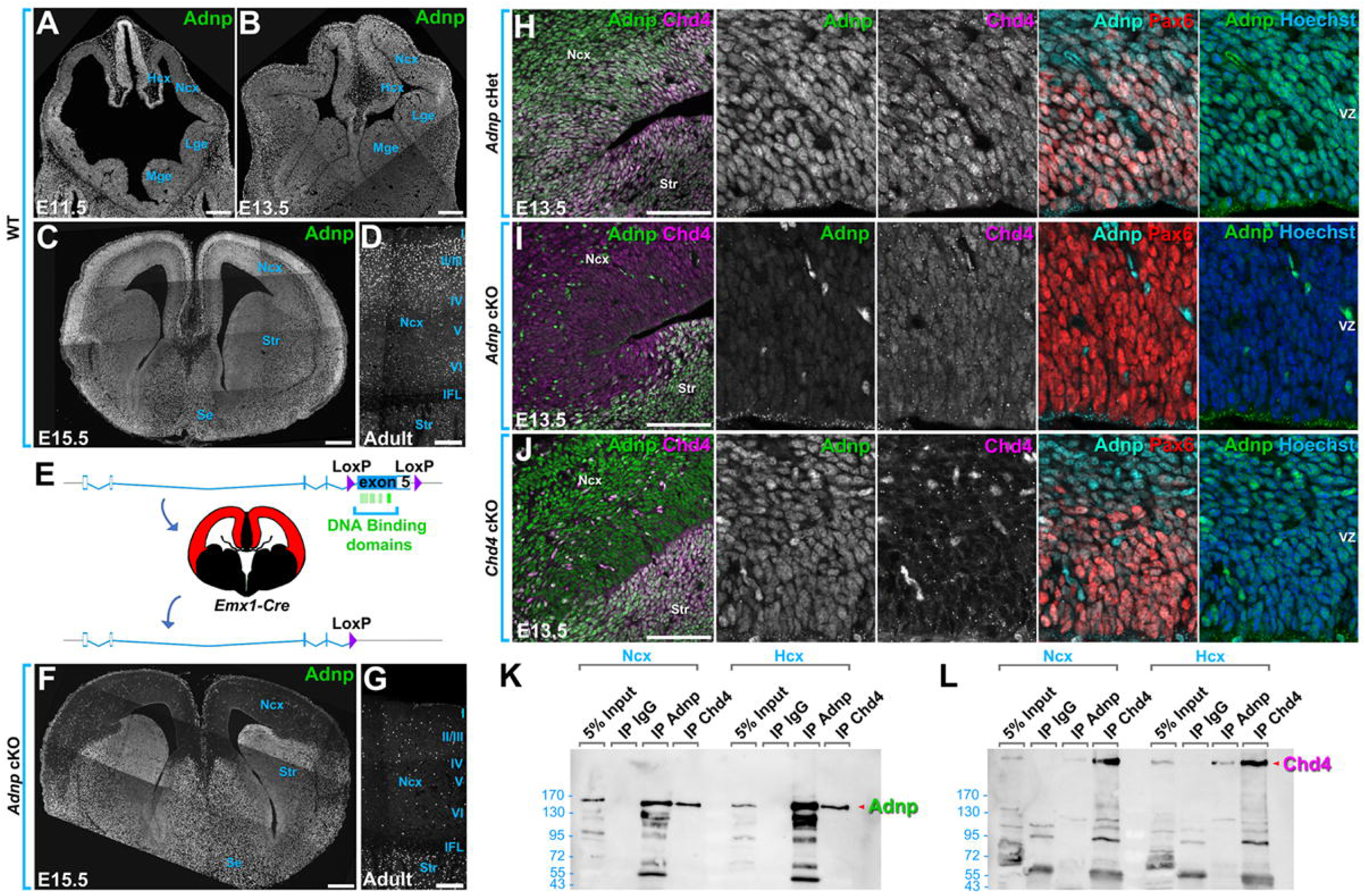
Conditional ablation of *Adnp* and Chd4 during neocortical neurogenesis. (A-E) Adnp immunohistochemistry on wild-type (WT) mouse brains at E11.5 (A), E13.5 (B), E15.5 (C), or adult (D) stages. (E) Conditional genetics strategy. Cre-mediated excision of LoxP-flanked exon 5 leads to the deletion of all but ∼60 amino acids from the Adnp coding sequence, including all of the DNA binding domains (green bars). Red color indicates the *Emx1-Cre* expression domain. (F, G) Adnp immunohistochemistry on *Adnp* cKO brains at E15.5 (F) or adult (G) stages. (H-J) Adnp, Chd4, Pax6, and DNA (Hoechst) staining on E13.5 *Adnp* cHet (H), *Adnp* cKO (I), and *Chd4* cKO (J) neocortices. (K, L) Co-immunoprecipitations performed on P3 neocortical or hippocampal protein lysates using the indicated antibodies. (K) Adnp western blot. (L) Chd4 western blot. Ncx: neocortex; Hcx: hippocampus; GE: ganglion eminence; LGE: lateral ganglionic eminence; MGE: medial ganglionic eminence; IFL: inner fiber layer; Se: septum; Str: striatum; VZ: ventricular zone. Scale bars = 200 μm.

Since germline *Adnp* knockouts were shown to be lethal at head-fold stages prior to the onset of neocortical neurogenesis^29^, we sought to generate a conditional *Adnp* allele. All but 67 of Adnp’s 1108 total amino acids are encoded within exon 5, including all of the putative DNA binding motifs (Fig. 1E). We therefore used CRISPR to surround this exon with flanking LoxP sites. Successful recombination was confirmed by long-range PCR and Southern blotting (Fig. S2B). We next crossed *Adnp^Flox^* mice with the *Emx1-Cre* driver, which is expressed specifically in the dorsal telencephalon beginning at E10.5^36^ to generate wild-type (WT; *Adnp^+/+^* and/or Cre-negative), *Adnp* conditional heterozygote (cHet; *Adnp^Flox/+^; Emx1-Cre+*), or *Adnp* conditional knockout (cKO; *Adnp^Flox/Flox^; Emx1-Cre+*) mice. *Adnp* cKOs were born in expected Mendelian ratios, and were viable and fertile. In accordance with expectations, Adnp protein expression in the cKO was abolished in the vast majority of cells within the dorsal telencephalon (Fig. 1F, G, I; Fig. S1). Within the neocortex, residual protein expression matched the expected pattern for cortical interneurons, which are outside of the *Emx1-Cre* lineage^36^ (see also Fig. 3H-J). Importantly, the loss of Adnp staining specifically in the dorsal telencephalon validates the knockout strategy, as well as the specificity of the Adnp antibody.

We additionally generated cKOs for the core ChAHP subunit *Chd4*, which is Adnp’s most quantitatively significant co-factor^2,14^. We utilized a *Chd4* conditional allele in which loxP sites flank exons 12-21, encompassing the ATPase domain^22^. Again, Chd4 protein was efficiently ablated from most cells within the dorsal telencephalon (Fig. 1J). In co-staining experiments with knockout-validated antibodies, we found that Adnp and Chd4 were broadly co-localized within the nuclei of virtually every cell, including Pax6+ apical progenitors (Fig. 1H-J; Fig. S1). To determine if the observed colocalization between Adnp and Chd4 proteins reflected *bona fide* protein-protein interactions we performed coimmunoprecipitations. When Adnp was immunoprecipitated from microdissected neocortical or hippocampal protein lysates, it readily pulled down Chd4. Likewise, Chd4 co-immunoprecipitated Adnp (Fig. 1K, L). Together, these data confirm the presence of the ChAHP complex within the developing neocortex.

Next, we examined ChAHP mutant brains. *Adnp* cKOs exhibited a gross reduction in the size of the dorsal telencephalon (Fig. 2A-C), which was evident via measurement of the 2 dimensional area of the dorsal telencephalon in wholemount images (Fig. 2G, H). Size differences were observed in both male and female cKOs (Fig. 2H). The reduced size of *Adnp* cKOs was evident as early as P2 and was not progressive over time (Fig. 2G), suggesting a prenatal or perinatal origin. With respect to *Chd4*, the Riccio lab previously reported perinatal lethality when *Chd4* cKOs were generated using the *Nestin-Cre* driver^13^. However, we recently reported that *Chd4* cKOs were viable when generated using *Emx1-Cre*, albeit at well below Mendelian ratios^37^. *Chd4* cKOs additionally exhibited sporadic hydrocephalus^37^ (Fig. 2F). At P28, *Adnp* and *Chd4* cKOs exhibited an indistinguishable ∼20% reduction versus WT or cHets when measured either as a function of cortical area or brain weight (Fig. 2I, J). These data are consistent with the notion that the ChAHP complex regulates cortical growth.

**Figure 2.**
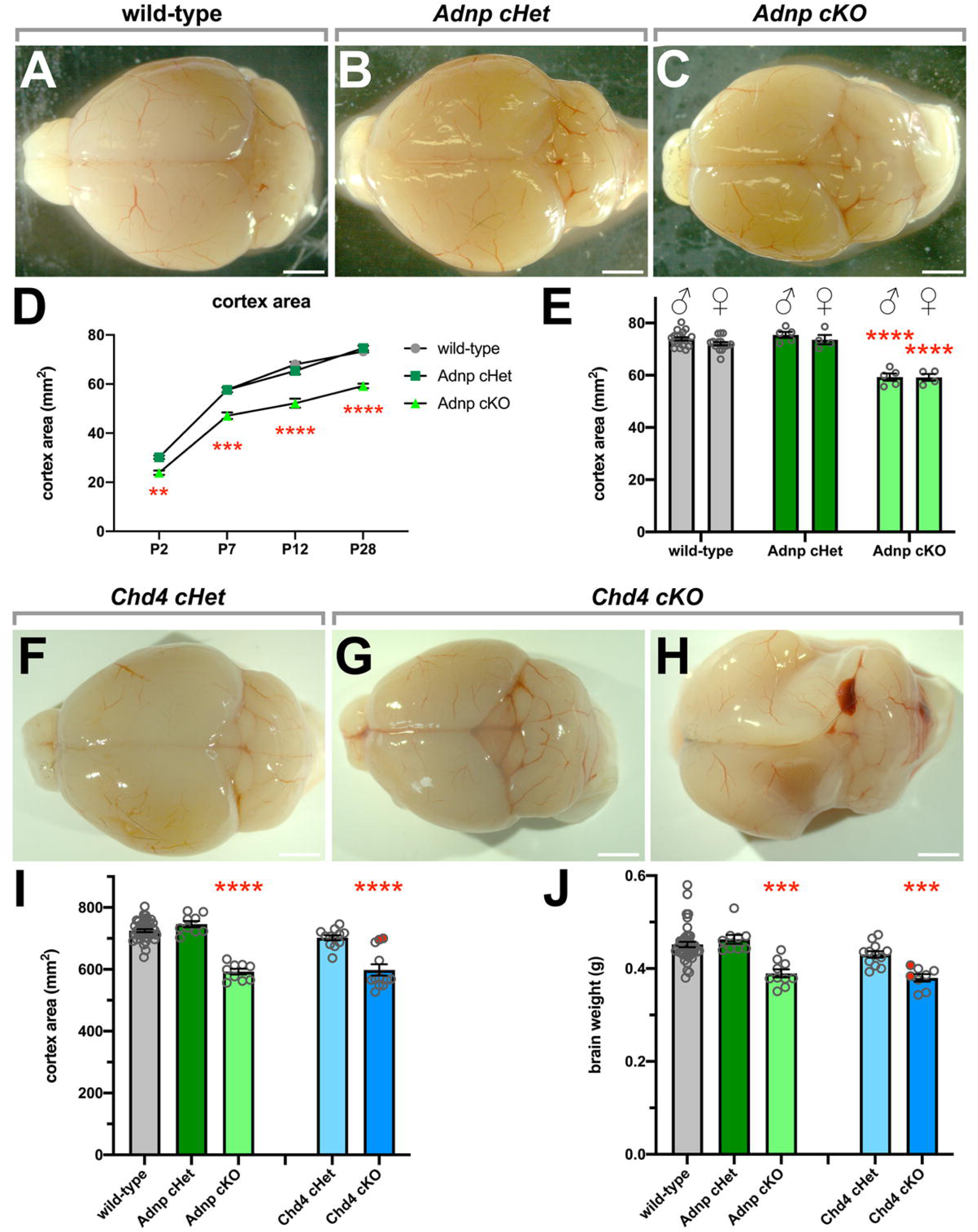
*Adnp* and *Chd4* are required for cortical growth. (A-F) Wild-type (A) *Adnp* cHet (B) *Adnp* cKO (C) brains harvested at P28. (D) Quantitation of cortex area at P2, P7, P12, and P28. n-values: P2: 6 cHet, 5 cKO; P7: 9 wt, 7 cHet, 3 cKO; P12: 9 WT, 5 cHet, 4 cKO; P28: 34 wt, 9 cHet, 9 cKO. ** p < 0.01 by Student’s t-test. *** p < 0.001, **** p < 0.0001 by one-way ANOVA with Tukey’s post-hoc test. (E) Quantitation of cortex area at P28 according to biological sex. **** p < 0.0001 by one-way ANOVA with Tukey’s post-hoc test comparing sexes separately. (F-H) *Chd4* cHet (F) and *Chd4* cKO (G, H) brains harvested at P28. The *Chd4* cKO in (H) is hydrocephalic. (I, J) Comparison of *Adnp* and *Chd4* cHet and cKOs by cortical area (I) or brain weight (J) at P28. Red data points indicate hydrocephalic brains. ** p < 0.01, *** p < 0.001, **** p < 0.0001 by Kruskal-Wallis with Dunn’s post-hoc test. Scale bars = 200 μm.

**Figure 3.**
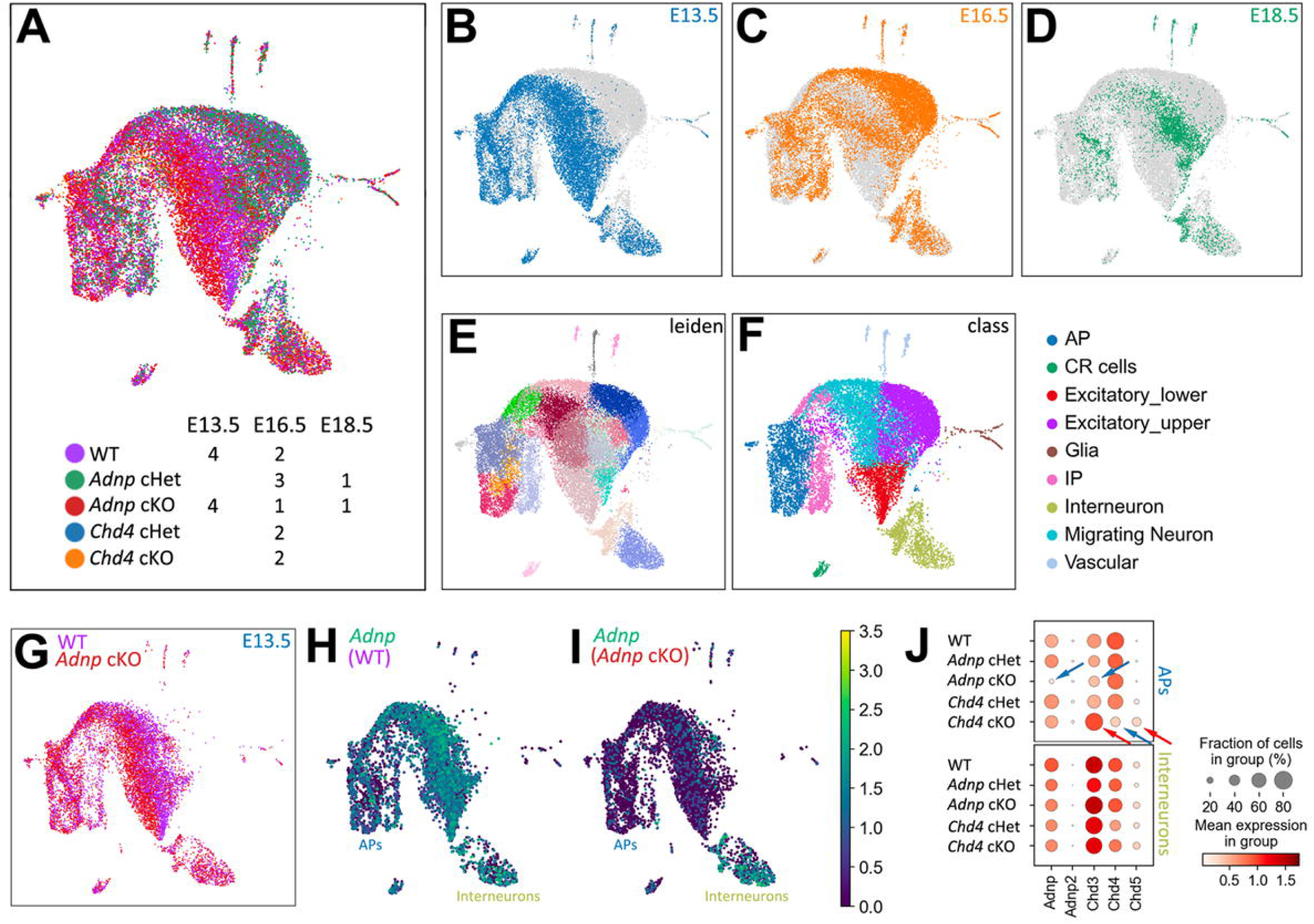
scRNA-seq atlas of neocortical development in *Adnp* and *Chd4* mutants. (A) UMAP embedding of 28,898 neocortical cells derived from the indicated genotypes and timepoints. (B-D) Individual timepoints as indicated. (E) Cluster identification via the Leiden algorithm. (F) Cell-types were annotated according to marker genes identified in a previous study^40^. See also Figs. S3 and S4. (G) Comparison of WT (n=4) and *Adnp* cKO (n=4) replicates at E13.5. (H, I) *Adnp* transcript expression in E13.5 WT (H) or *Adnp* cKO (I) replicates. (J) Comparison of *Adnp* and *Chd4* paralog gene expression from all timepoints in APs (top) or Interneurons (bottom). Note that interneurons fall outside of the *Emx1-Cre* lineage, and therefore serve as an internal control. Blue arrows: downregulated in *Adnp* cKO; Red arrows: upregulated in *Adnp* cKO. AP: apical progenitors; IP: intermediate progenitors; CR: Cajal-Retzius cells.

### Adnp and Chd4 program developmental trajectories during neocortical development

Since ChAHP has been shown to regulate gene expression and cell fate decisions^2,20,29,30^, we hypothesized that *Adnp* and *Chd4* cKOs might exhibit similarly dysregulated gene expression programs, which could be global or restricted to specific cell subtypes. To characterize the developmental trajectories of neocortical cells in a comprehensive fashion we obtained single-cell (sc) transcriptomic profiles of neocortices from *Adnp* and *Chd4* mutants. We profiled mice at E13.5, E16.5, and E18.5. At E13.5, the neocortex is mainly producing deep-layer (VI and V) neurons, whereas at E16.5, it almost exclusively produces upper-layer (II/III) neurons^38^. At E18.5, neurogenesis ceases, and progenitor cells switch to producing glia. In total, we profiled 28,898 high-quality cells (Methods). In these experiments, we used the Multi-seq^39^ approach, which relies on lipid-tagged barcodes in order to facilitate multiplexing of samples prior to scRNA-seq. Samples from different embryos were individually barcoded, combined, and then subjected to conventional scRNA-seq using the 10X Genomics Chromium platform. We profiled 6 WT, 4 *Adnp* cHet, and 6 *Adnp* cKO embryos, as well as 2 *Chd4* cHet and 2 *Chd4* cKO neocortices (Fig. 3A-D, Table S1).

Gene expression matrices from different samples were combined, following which, standard procedures of normalization, batch correction, and dimensionality reduction were applied (Methods). Clusters of transcriptomically related cells were determined in the reduced dimensional space using the Leiden algorithm (Fig. 3E). We next annotated each cluster into cortical cell types based on the expression of marker genes defined in a previous study of the developing neocortex^40^ (Fig. S3C-H; Fig. S3, S4). The relative arrangement of the subclasses in UMAP space mirrored the known developmental progression from apical progenitors to intermediate progenitors and/or migrating neurons, and finally to lower layer or upper layer neurons. Within this scaffold, *Adnp* WT, cHet and cKO cells largely overlapped, although some segregation of cKOs from other samples was evident (Fig. 3G), which was substantiated by statistically significant transcriptional differences (see below).

As expected, visualization of *Adnp* transcripts revealed an almost complete loss of expression in *Adnp* cKO cells, except for interneurons, which are outside of the *Emx1-Cre* lineage^36^ (Fig. 3H, I). In *Chd4* cKOs, *Chd4* transcription was reduced in apical progenitors, but not eliminated. This reflects the fact that the genetic strategy excises the ATPase/helicase domain but leaves the 3′ region of the gene intact and in-frame^41^. Importantly, while residual *Chd4* transcript was detected in the *Chd4* cKO, Chd4 protein was not detected, indicating that recombination of the loxp-flanked cassette was efficient (Fig. 1J). We next examined the expression of ChAHP subunit paralogs (ie. *Adnp/Adnp2* and *Chd3/4/5*). In *Adnp* cKOs, compensatory upregulation of the paralogous *Adnp2* gene was not detected (Fig. 3J). We observed that both *Chd3* and *Chd5* were upregulated in *Chd4* cKO cells, suggesting possible genetic compensation. We did not detect cross-regulation between *Adnp* and *Chd4*, as *Chd4* levels were unaffected in the *Adnp* cKO and *vice versa* (Fig. 3J). However, surprisingly, we saw a *reduction* in *Chd3* expression in the *Adnp* cKO (Fig. 3J; see also below). Finally, gene expression was unaffected in interneurons in accordance with the lack of Cre expression in this lineage. Taken together, these data collectively validate the genetic models as well the gene expression data.

### Adnp is required for the expansion of upper neocortical layers

Since neurodevelopmental phenotypes have been previously characterized in the *Chd4* cKO^13,26^, we focused on the *Adnp* cKO. We first examined differentially expressed genes (DEGs) in mutant scRNA-seq samples. To avoid batch effects, we made pairwise comparisons between mutants and controls that were processed together, rather than in separate experiments. We found that there was good overlap between DEGs identified from all 3 timepoints (Fig. 4A; Supplemental data File 1). E13.5 *Adnp* cKO samples had the most DEGs (1291 genes, adj. p-value <0.05), consistent with the fact that the E13.5 experiment was the best-powered. However, the majority of DEGs identified at later timepoints were also differential at E13.5 (Fig. 4A). Results using an alternate DEG identification method^42^ were highly consistent (Supplemental data file 2). To examine heterozygotes, we compared WT and *Adnp* cHets at E16.5. However, only 18 DEGs were identified, the majority of which were also differentially regulated in *Adnp* cKOs (Fig. 4A; Supplemental data File 1). As an internal control, we also examined cortical interneurons, which were captured alongside cKO cells, but are outside of the *Emx1-Cre* lineage. In these cells, the only DEGs identified were *Xist* and *Tsix*, which are transcribed from inactivated X chromosomes and thus likely reflect sex imbalances between the selected cKO and control embryos (see Fig. S5).

**Figure 4.**
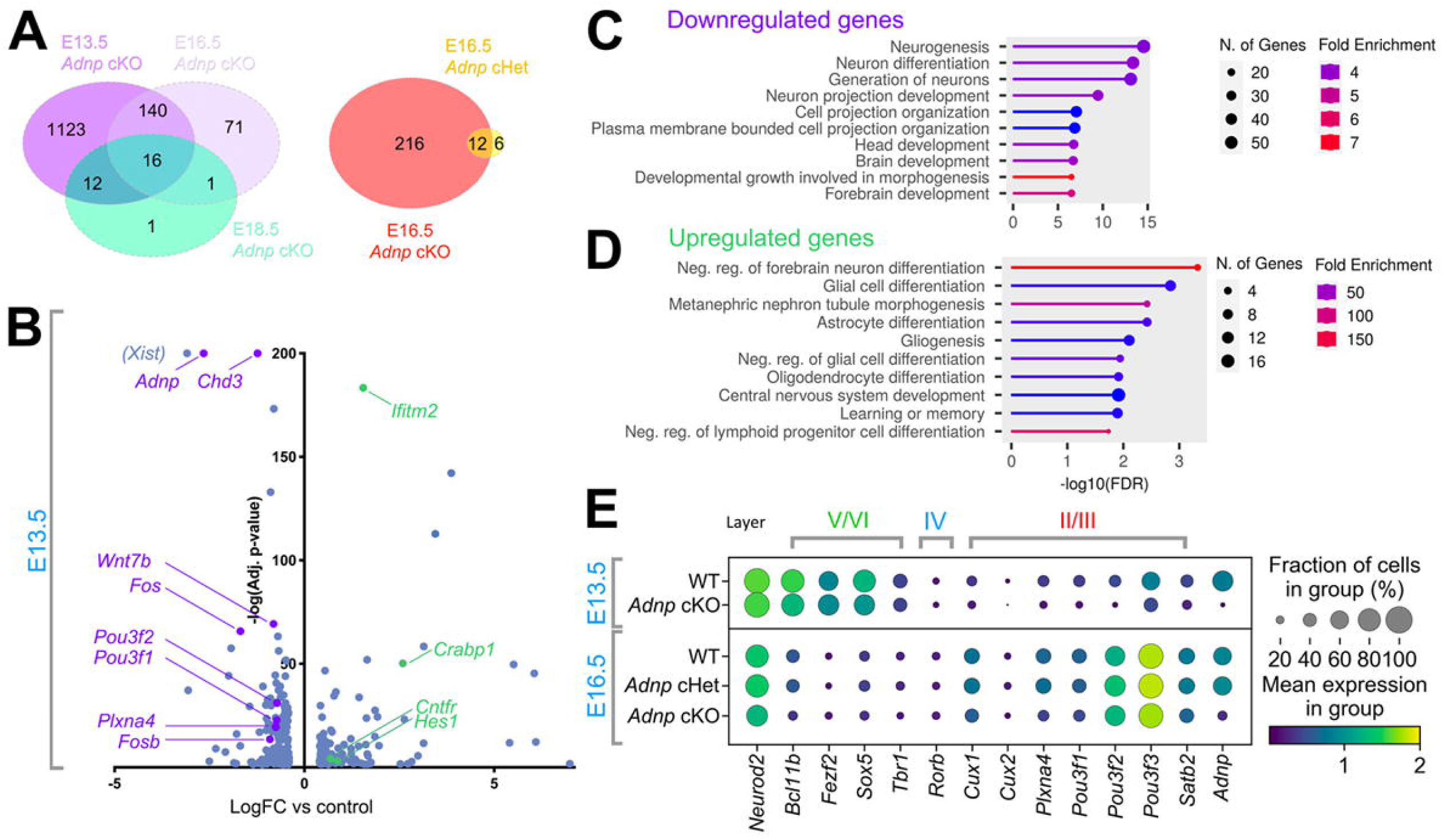
*Adnp* regulates neurogenic gene expression. (A) Venn diagrams of overlap between differentially expressed genes (adj. p-value <0.05) in *Adnp* cKO at E13.5, E16.5, and E18.5, or in E16.5 *Adnp* cHet versus cKO. (B) Volcano plot of DEGs (adj. p-value <0.05; LogFC > 0.4 or < −0.4) from *Adnp* cKO samples vs. wild-type control at E13.5. Selected significantly downregulated genes are depicted in purple, while upregulated genes are depicted in green. The *Xist* gene reflects imbalances in biological sex (see also Fig. S5). (B, C) Top 10 GO terms (by FDR and LogFC) for *Biological Process* on genes depicted in (A). (E) Expression of neuronal markers in E13.5 and E16.5 excitatory neurons (lower-layer and upper-layer). Roman numerals indicate neocortical layers.

At E13.5, *Adnp* cKOs exhibited both downregulated and upregulated genes (Fig. 4B). Interestingly, several upper-layer marker genes were significantly downregulated, including *Plxna4* (logFC = −0.75), *Pou3f1* (logFC = −0.73), and *Pou3f2* (logFC = −0.72; Fig. 4B). Upregulated genes included *Crabp1* (logFC = 2.6), *Cntfr* (logFC = 0.69), *Hes1* (logFC = 0.87), and *Ifitm2* (logFC = 1.55) – all of which are glial markers or determinants. Gene ontology analysis revealed that as a group, the downregulated genes were associated with neurogenesis and cortical growth, whereas the upregulated genes were associated with gliogenesis, or with negative regulation of neurogenesis (Fig. 4C, D). Examining gene expression specifically in excitatory neurons, we found that lower-layer marker genes were not markedly dysregulated, but that upper-layer marker genes were systematically downregulated – both at E13.5 and E16.5 (Fig. 4E).

To confirm these observations, we examined the *Adnp* cKO neocortex histologically at E15.5, during the peak of upper-layer neuron production. We first measured the overall thickness of the neocortex and counted the total numbers of Hoechst+ nuclei. The gross reduction in cortical size observed postnatally was not yet statistically significant at E15.5 (Fig. 5A, B, K). Since WT and cHet animals also exhibited little difference in gene expression (Fig. 3A), we therefore pooled these genotypes together as ‘controls’ for subsequent analyses. We next visualized neurons using markers of early-born (Bcl11b/Ctip2) and late-born (Pou3f2/Brn2) subtypes. Again, counting in 100 micron-wide bins, we found no difference in absolute or proportional numbers of Bcl11b+ cells (Fig. 5A-D, L). Pou3f2+ neurons were contrastingly reduced by ∼25% in the *Adnp* cKO CP (Fig. 5E-J, M). In addition to CP neurons, Pou3f2 also marks cells in the VZ, SVZ, and IZ. However, in the *Adnp* cKO, Pou3f2 levels were unaffected in these zones (Fig. 5M), suggesting that the defect in Pou3f2 expression was restricted to neurons that had reached the CP.

**Figure 5.**
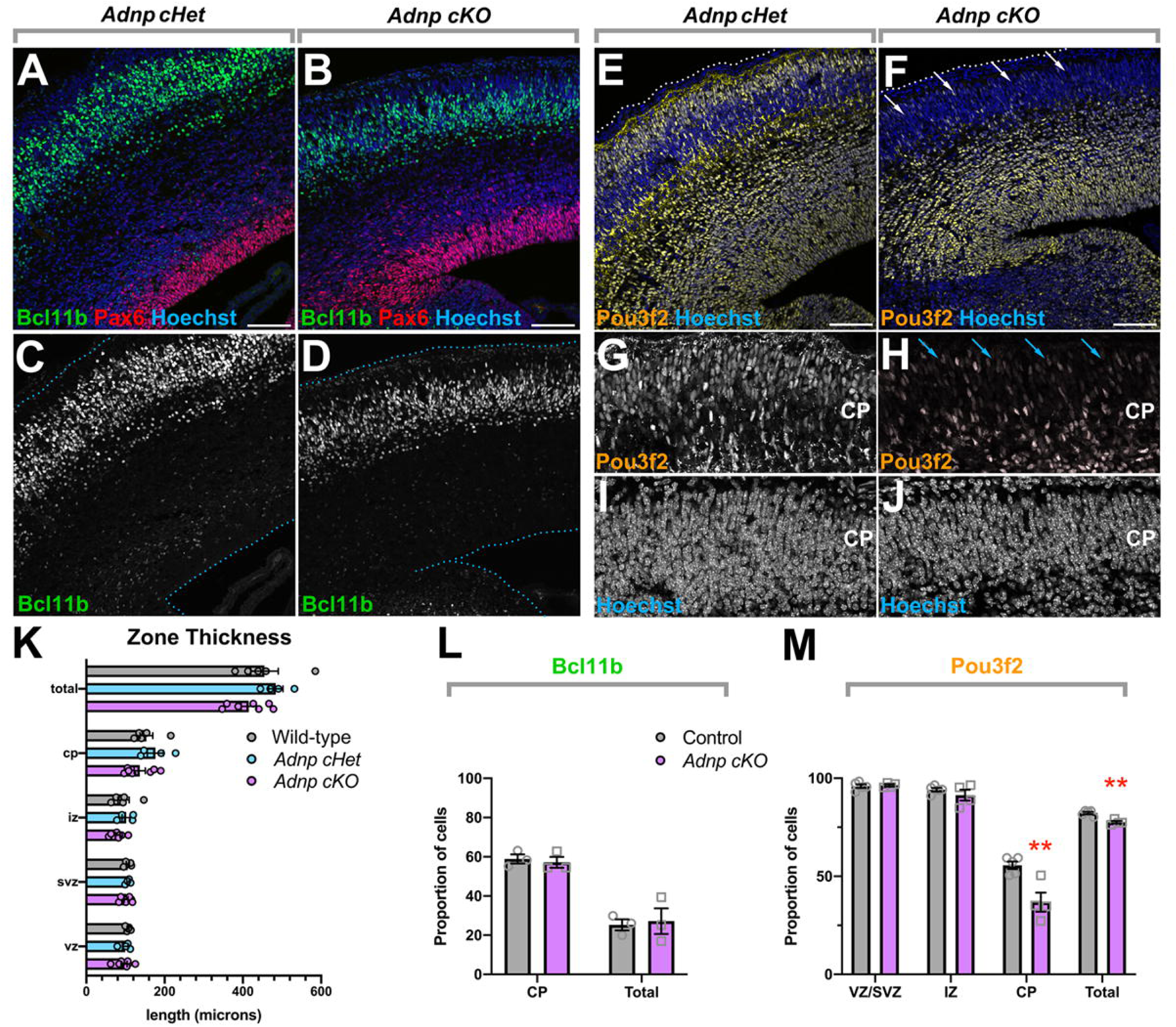
*Adnp* is required for the production of upper-layer neurons. (A-J) Immunohistochemistry for cell-type markers in the E15.5 embryonic neocortex. (A-D) Immunohistochemistry for the apical progenitor marker Pax6 and the lower-layer marker Bcl11b. (E-J) Immunohistochemistry for Pou3f2. Pou3f2 marks progenitors and differentiating neuronal precursors, as well as upper-layer neurons that have migrated to the cortical plate. This latter population is reduced in the *Adnp* cKO (G-J; arrows). (K) Measurement of the size of neocortical zones in wild-type, cHet, and, cKO brains. (L) Quantitation of Bcl11b levels by zone. (M) Quantitation of Pou3f2 levels by zone. ** p < 0.01 via two-way Student’s t-test. VZ: ventricular zone; SVZ: subventricular zone; CP: cortical plate. Scale bars = 200 μm.

Since neurogenesis is not complete at E15.5, we also examined mature brains at P28. In cross-section, we noted that the overall thickness of the cortical plate was reduced in *Adnp* cKOs in accordance with wholemount measurements (Fig. S6A-E). We counted cells in 200 micron wide bins, revealing that total cell numbers were reduced by approximately 20% in *Adnp* cKOs versus controls, although the density of cells per square micron was unaffected (Fig. S6E). Similar to E15.5 counts, we found that the overall number of Bcl11b+ cells was not different between control and *Adnp* cKO brains. However, Pou3f2+ cells were strikingly reduced in the *Adnp* cKO versus controls (Fig. S6F). The expression of other layer-biased marker proteins, such as Tbr1, and Satb2 similarly suggested that although neurons of all layers were present in the *Adnp* cKO, the size of the upper-layers was specifically reduced. These data confirm that the underproduction of upper-layer neurons contributes to the hypoplasia observed in *Adnp* cKOs. This phenotype is strikingly similar to that reported for *Chd4* cKOs, in which upper-layer neurons are also specifically underproduced^13,26^.

### Adnp is required for apical progenitor proliferation during late phases of neurogenesis

To pinpoint the source of the observed reduction of upper-layer neurons, we first examined cell death at 2 embryonic timepoints. We performed immunohistochemistry for active caspase 3, and quantitated pyknotic nuclei from tissue sections, but observed only trace levels of cell death in both controls and cKOs (Fig S7).

To examine progenitor proliferation and self-renewal, we marked S-phase cells with EdU and harvested embryos 24 h later, from E12.5→E13.5 (Fig. 6A-D) or E14.5→E15.5 (Fig. 6E-N). Embryos were sectioned and stained for Ki67, which marks all proliferating cells. At E13.5, we observed no difference between EdU or Ki67 labelling in control (Cre-negative) brains vs. *Adnp* cKOs (Fig. 6A, B). Most cells within the VZ were Ki67+, and almost all EdU+ cells were accordingly double-positive (Fig. 6C). Within the SVZ/CP, cells were more frequently marker-negative, but we observed no difference between *Adnp* cKOs and controls (Fig. 6D). However, at E15.5, control brains exhibited a band of EdU+ nuclei at the apical surface (Fig. 6E). Most of these cells were doubly positive for Ki67, indicating that the cells had passed through S phase and continued to proliferate over the 24 h interval. A second band of EdU+ cells localized at the basal margin of the VZ. Many of these cells were Ki67-negative, indicating that over the course of 24 h, they had undergone cell cycle exit (Fig. 6E). In *Adnp* cKOs, while the band of EdU+ cells at the basal margin of the VZ was retained, EdU labelling was dramatically reduced from the apical surface of the VZ (Fig. 6F; arrows). Accordingly, the overall proportions of Ki67+ or EdU+ cells within the VZ were reduced by ∼25% (Fig. 6G). Moreover, Ki67/EdU double-positive cells were reduced both in absolute terms, and relative to EdU single positive cells (Fig. 6G), suggesting deficits in self-renewal specifically during the developmental window for upper-layer neuron production.

**Figure 6.**
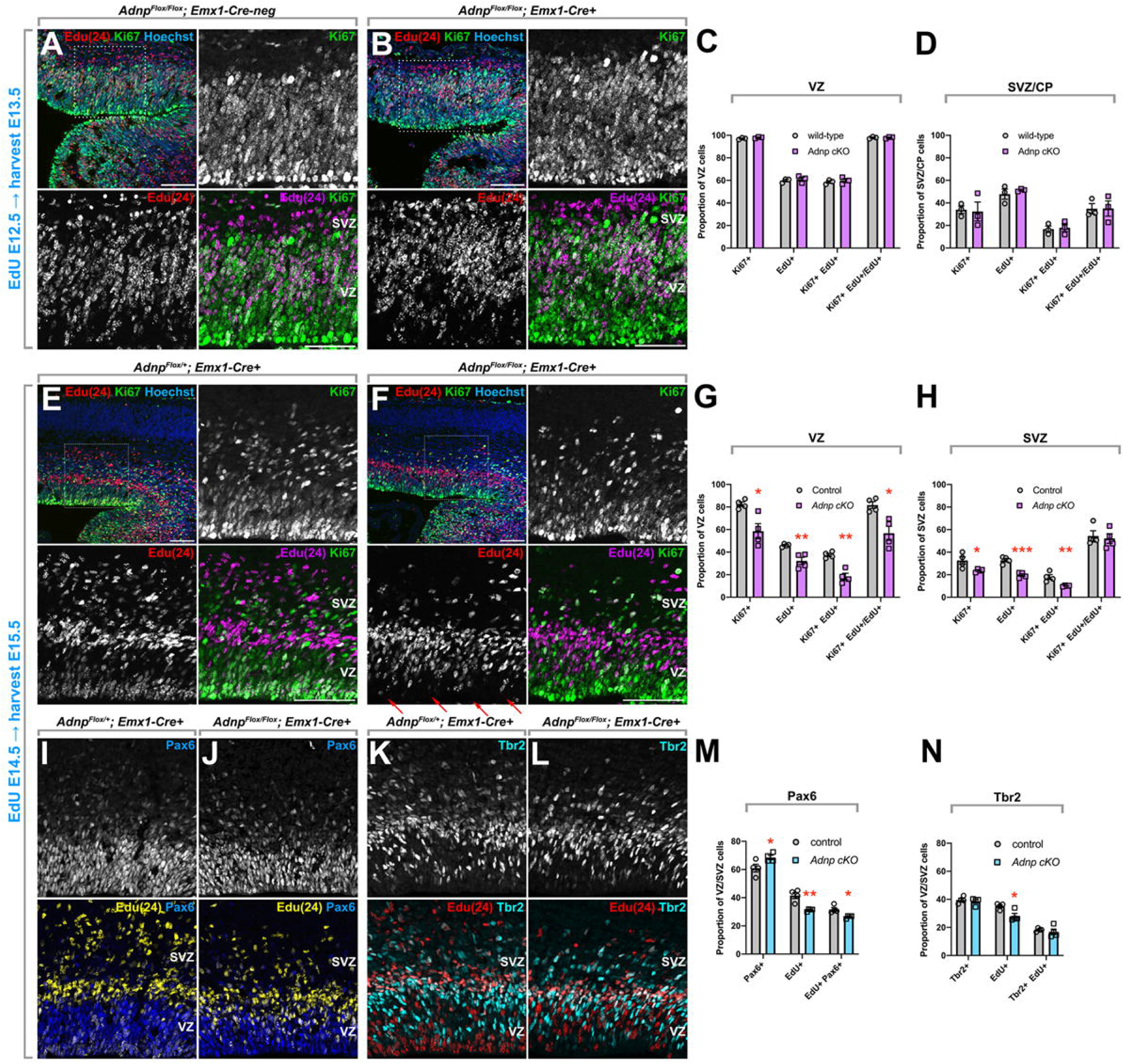
Adnp is required for apical progenitor proliferation and self-renewal. EdU was administered 24 hours prior to harvest at E13.5 (A-D) or E15.5 (E-N). Upper left panels in (A, B, E, F) depict the regions shown at higher magnification in the other panels (dotted squares). (C, D, G, H) Quantitation of the proportion of cells that are EdU+, Ki67+, double-positive, or the fraction of double-positive cells among total EdU+ cells within the ventricular zone (C, G) or subventricular zone/cortical plate (D, H) as indicated. (I-N) At E15.5, EdU was co-stained with the apical progenitor marker Pax6 (I, J) or the basal progenitor marker Tbr2 (K, L). (M) Quantitation of the proportion of EdU+, Pax6+, or double-positive cells within the VZ and SVZ. (N) Quantitation of the proportion of EdU+, Tbr2+, or double-positive cells within the VZ and SVZ. * p < 0.05; ** p < 0.01; *** p < 0.001 by Student’s t-test. VZ: ventricular zone; SVZ: subventricular zone. Scale bars = 100 μm.

We next examined the SVZ. At E15.5, although the overall proportion of cells that were marked by EdU or Ki67 was lower in the SVZ versus the VZ, we observed a similar reduction in EdU/Ki67 single positive cell proportions, but the rate of EdU/Ki67 double-positive cells among total EdU+ cells was not different between the genotypes, suggesting that progenitor self-renewal in the SVZ was unaffected.

Since the VZ is enriched in apical progenitors whereas the SVZ is enriched in basal progenitors, we sought to visualize these populations specifically using Pax6 or Tbr2 immunohistochemistry, respectively (Fig. 6I-L) - this time counting cells irrespective of the VZ/SVZ boundary in order to avoid potential bias in the assignment of these zones. In accordance with our VZ/SVZ Ki67 counts, we found that the overall proportions of EdU+/Pax6+ apical progenitors were reduced in *Adnp* cKOs (Fig. 6I, J, M), but there was no difference in the overall numbers of Tbr2+ or Tbr2/EdU double-positive cells (Fig. 6K, L, N). Thus, while Pax6+ apical progenitors were not decreased in numbers, our data suggests that *Adnp* is required for the proliferation and self-renewal of neocortical apical progenitors specifically at late-stages. These proliferative defects likely underlie the gross hypoplasia exhibited by the *Adnp* cKO.

### Adnp and Chd4 cooperate to regulate cortical growth

To confirm that ChAHP regulates the genome during cortical development, we next asked whether Adnp and Chd4 proteins occupy the same gene regulatory elements. While the genomic occupancy of Adnp during neocortical development was previously reported^18^, Chd4 genomic occupancy during cortical development had not yet been examined. We therefore performed cut&run-seq on Chd4 using an antibody previously validated in ChIP-seq^27^. We compared these data with ChIP-seq tracks for histone modifications previously generated by the Bing Ren laboratory^43^. Since ChAHP had been reported to compete with the Ctcf transcription factor for genomic occupancy^15^, we also generated Ctcf cut&run-seq data for comparison. All of the above cut&run-seq datasets were generated from E13.5 murine cortices, while the ChIP-seq datasets were generated from E13.5 forebrain samples. Examining H3K4me3, we found that approximately half of the Chd4 cut&run peaks mapped to active promoters (Fig. 7A: cluster C1), while the other half mapped to enhancer loci co-occupied by Ctcf and/or H3K27ac but devoid of H3K4me3 (cluster C2). Adnp cut&run signal was mainly found at Chd4-occupied promoters, suggesting a role for ChAHP in gene regulation.

**Figure 7.**
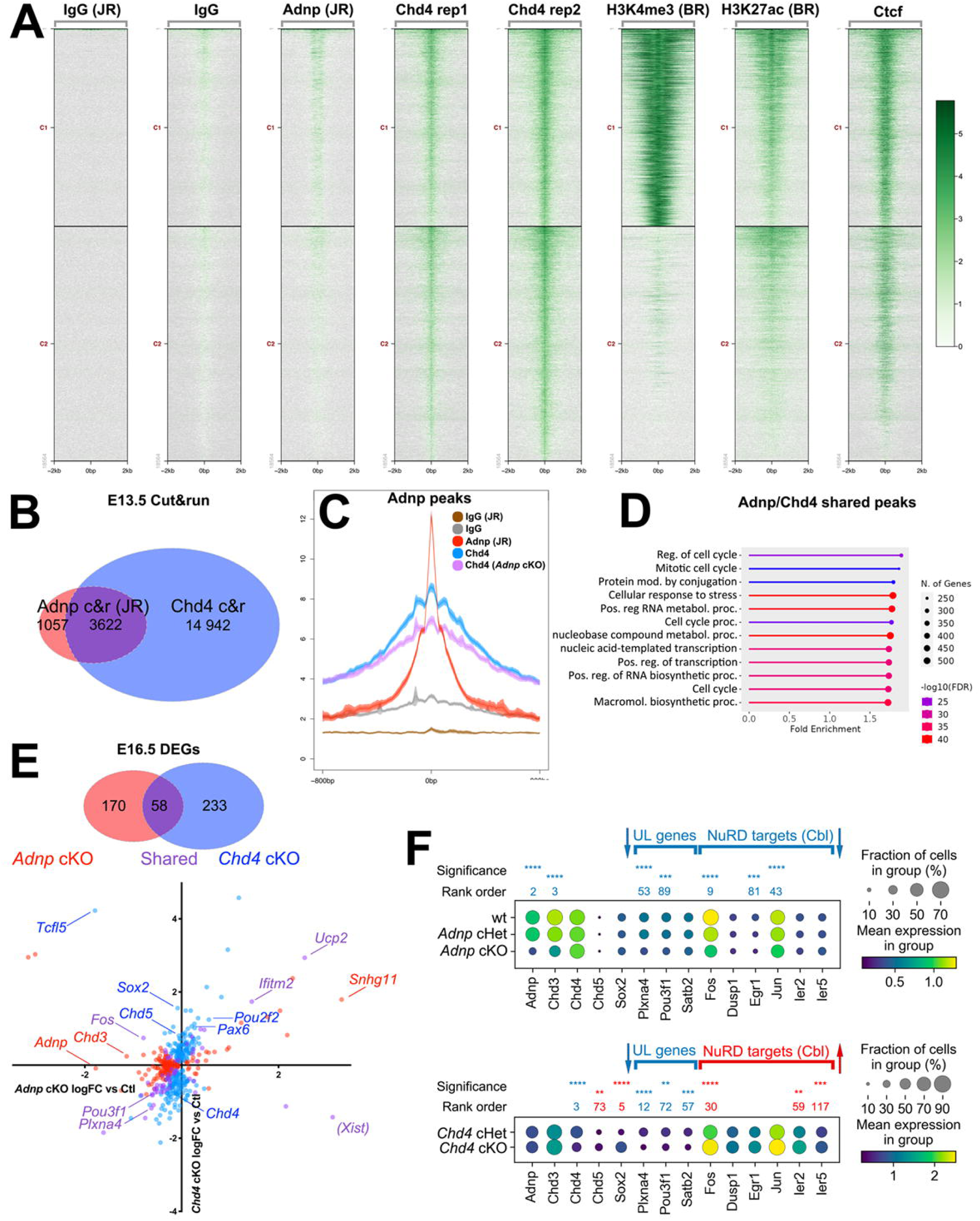
(A) Cut&run-seq comparing the genomic occupancy of Chd4 or Ctcf, with an Adnp dataset previously generated by the John Rubinstein lab (JR)^18^ – all from E13.5 forebrain/cortical tissue. Datasets were also compared against ChIP-seq for H3K4me3 (marking active promoters) and H3K27ac (marking promoter/enhancer elements) generated by the Bing Ren lab (BR) from E13.5 forebrain^43^. Data are centered on (wild-type) Chd4 peak summits. (B) Venn diagram of called peaks for Chd4 (blue) or Adnp (red). (C) Comparison of Chd4 occupancy in wild-type (blue) or *Adnp* cKO (purple) E13.5 cortices at Adnp peak loci. (D) GO terms analysis for Biological Function on genes co-occupied by Adnp and Chd4. (E) Venn diagram of overlap between differentially expressed genes in *Adnp* cKO vs. control (red), or *Chd4* cKO vs. control (blue). Shared genes are depicted in purple. (F) Scatterplot of differentially expressed genes comparing *Adnp* cKO vs. WT (X-axis), or *Chd4* cKO vs. cHet control (Y-axis). (G) Regulation of previously identified Chd4 target genes^28^ at E16.5. Dotplots depict gene expression in all cells. Blue text/arrows: significantly downregulated. Red text/arrows: significantly upregulated. * p < 0.05, ** p < 0.01, *** p < 0.001, **** p < 0.0001 by adjusted p-value (Wilcoxon rank sum test).

Chd4 bound considerably more loci in comparison with Adnp. Using MACS2^44^ to call peaks on both datasets, we found that Chd4 occupied 18 564 genomic loci, whereas the published Adnp cut&run-seq dataset yielded 4679 peaks. 80% of Adnp peaks were co-occupied by Chd4 (Fig. 7B). In the *Adnp* cKO, Chd4 signal was reduced at Adnp peak locations (Fig. 7C), suggesting that Adnp is required to recruit ChAHP to the genome. At loci co-occupied by Adnp and Chd4, peak-to-gene annotation followed by GO terms analysis for Biological Function revealed a significant enrichment of terms related to proliferation (Fig. 7D).

We additionally observed that the majority of DEGs identified in our E13.5 *Adnp* cKO transcriptomic data were occupied by Adnp and/or Chd4 (Fig. S8A). Co-occupied DEGs included *Cdk4*, *Ctnnb1*, *Hes5*, *Neurog2*, and *Pou3f2* – all of which regulate progenitor proliferation (Fig. S8B-K). Systematic analysis of cell cycle-dependent gene expression did not reveal marked changes at E13.5 (Fig. S8L, M), in accordance with the lack of proliferative defects observed in histological experiments (see Fig. 6). However, the expression of many proliferation genes was reduced in the *Adnp* cKO (Fig. S8N) – perhaps prefiguring the deficits exhibited by apical progenitors at E15.5. Thus, integration of E13.5 cut&run-seq data with DEGs identified in scRNA-seq data suggests that ChAHP directly regulates genes that control progenitor proliferation and neurogenesis.

### Coordinate and divergent gene regulation by Adnp and Chd4

We next examined our E16.5 scRNA-seq dataset in order to directly compare the requirement for Adnp versus Chd4 for gene regulation. We examined DEGs (Adj. p-value < 0.05) identified in *Adnp* or *Chd4* cKOs versus their respective control samples. We found that a substantial fraction of these DEGs were shared between *Adnp* and *Chd4* cKOs (Fig. 7E; Supplemental datafile 1). Systematic comparison of *Adnp* versus *Chd4* DEGs revealed a weak positive correlation. However, while DEGs from *Chd4* cKOs were both upregulated and downregulated, most Adnp DEGs were downregulated at E16.5 (Fig. 7E). Thus, some transcripts were coordinately regulated by Adnp and Chd4, while others exhibited divergent regulation.

Next, we examined specific marker genes. A previous study had shown that Sox2 was significantly upregulated in *Chd4* cKO neocortical progenitors at E16.5^13^. We found that Sox2 was accordingly upregulated in *Chd4* cKOs scRNA-seq data (Fig. 7F). However, *Sox2* expression was not affected in *Adnp* cKOs. In accordance with the defective production of upper-layer neurons common to both *Adnp* and *Chd4* cKOs^13,26^, we observed that upper-layer markers such as *Plxna4*, *Pou3f1*, and *Satb2* were significantly downregulated in both *Adnp* and *Chd4* mutants (Fig. 7F), consistent with the idea that these genes might be regulated by ChAHP.

By contrast, we observed that some genes were regulated oppositely by Adnp and Chd4. In the cerebellum, Chd4 had been previously shown to suppress a variety of genes, including *Chd5*^27^, as well as the immediate early genes *Fos*, *Dusp1*, and *Jun*^28^, acting via the NuRD complex. Similarly, we found that many immediate early genes were differentially upregulated in the *Chd4* cKO neocortex. Surprisingly, these genes were oppositely regulated in the *Adnp* cKO (Fig. 7F). These latter observations are perhaps consistent with the hypothesis that some targets might be regulated oppositely by ChAHP versus NuRD.

### Adnp regulates a network of neurodevelopmental disorder genes

To gain insight into the relationship between Adnp and NDDs, we examined the SFARI gene database^45^, which curates human genes linked to ASD. We first intersected SFARI genes with E13.5 *Adnp* cKOs DEGs (Fig. 8A; Supplemental datafile 3). We found that of the 1291 DEGs identified in E13.5 *Adnp* cKOs, 126 were orthologues of SFARI genes. After excluding *Adnp* itself, we found that SFARI genes were enriched approximately 2.9-fold above expectations (p = 7.54−10^-27^, hypergeometric test). Next, we performed a similar intersection between DEGs and ‘definitive’ genes from the SysNDD database^46^, which curates genes associated with neurodevelopmental disorders, including ASD and ID. Again, we found that 172 DEGs were orthologous to definitive SysNDD genes. Excluding *Adnp*, this represents a 2.71-fold enrichment above expectations (p = 3.90−10^-33^, hypergeometric test). 74 DEGs overlapped between the SFARI and SysNDD lists (Fig. 8A, B). Next, we examined the 284 DEGs identified in *Chd4* cKOs at E16.5. We found that 44 DEGs matched the mouse orthologues of SFARI genes (4.52-fold enrichment vs. expectation, p = 6.10−10^-17^, hypergeometric test). 54 DEGs matched the SysNDD database (3.81-fold enrichment vs. expectation, p = 1.82−10^-17^ hypergeometric test, excluding *Chd4*). 29 genes matched both lists. Finally, we identified 20 genes common to all four groups, including *Celf2*, *Mef2c*, *Myt1l*, and *Satb2* (Fig. 8A-C).

**Figure 8.**
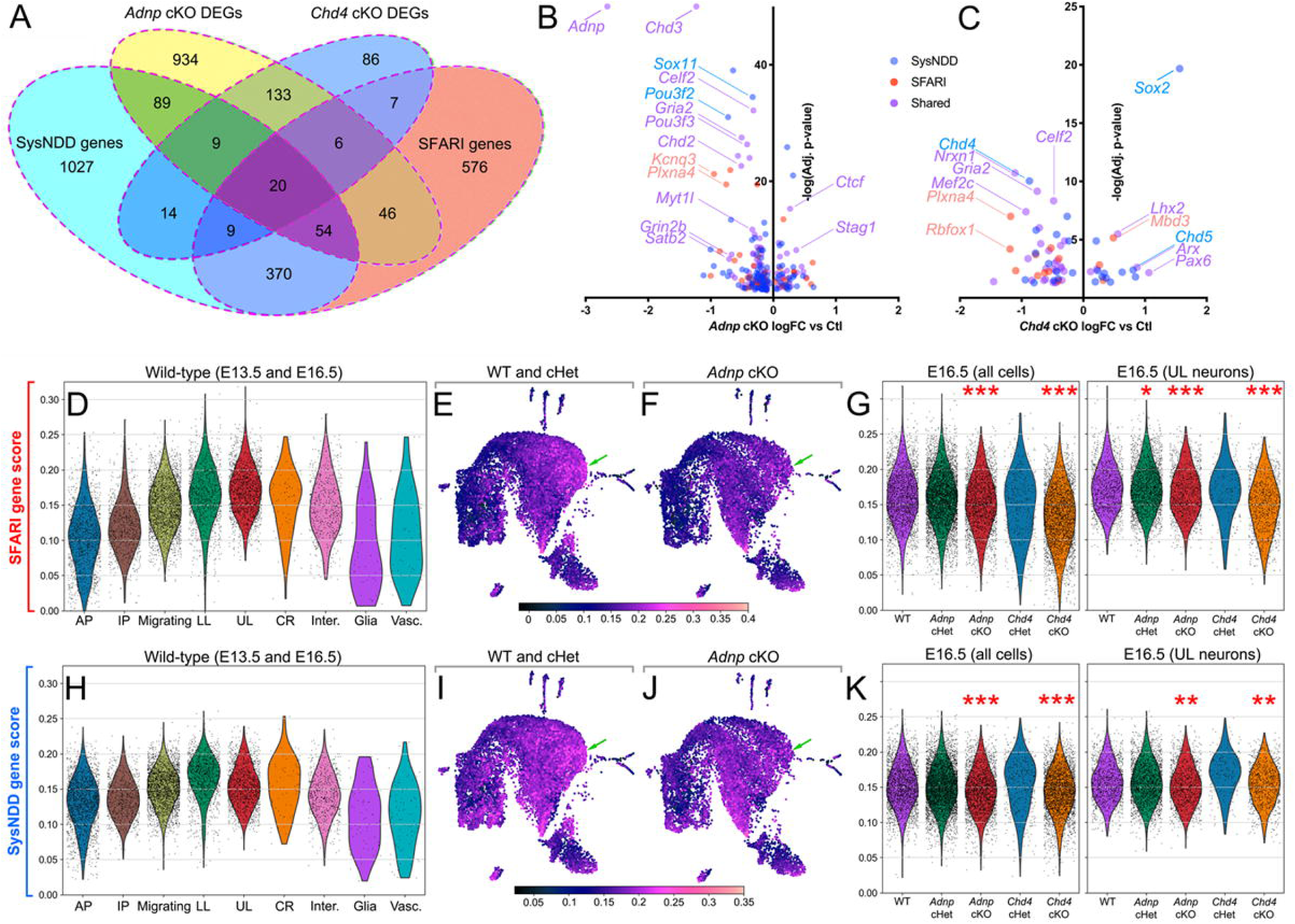
Adnp and Chd4 directly regulate neurodevelopmental disorder risk genes. (A) Venn diagram of mouse orthologues of SFARI ASD risk genes, SysNDD risk genes, E13.5 *Adnp* cKO DEGs, or E16.5 *Chd4* cKO DEGs. (B) Volcano plot of E13.5 *Adnp* cKO DEGs orthologous to SFARI genes (purple), SysNDD genes (blue), or both (red). (C) Volcano plot of E16.5 *Chd4* cKO DEGs orthologous to SFARI genes or SysNDD genes. (D) Violin plot of SFARI scores in WT cells (E13.5 and E16.5) according to cell type. (E, F) UMAP projections of E16.5 gene expression scores for mouse orthologues of SFARI risk genes in WT (E) or *Adnp* cKO (F). Arrows mark upper-layer neurons. (G) Violin plots of gene expression scores for mouse orthologues of SFARI risk genes in all cells, or upper layer neurons as indicated. (H) Violin plot of SysNDD scores in WT cells (E13.5 and E16.5) according to cell type. (I, J) UMAP projections of E16.5 gene expression scores for mouse orthologues of SysNDD risk genes in WT (I) or *Adnp* cKO (J). (K) Violin plots of gene expression scores for mouse orthologues of SFARI risk genes in all cells, or upper layer neurons as indicated. * p-value < 0.05, ** p-value < 0.01, *** p-value < 0.001 by ANOVA with Tukey’s post-hoc test.

We next obtained gene scores for SFARI or SysNDD risk genes, in which the average expression of SFARI or SysNDD risk genes was computed for each cell (normalized against the expression value for all genes). When SFARI gene scores were examined in E13.5 and E16.5 wild-type replicates, we noted that the expression of these genes was particularly prominent within neurons (Fig. 8D, E) – especially upper-layer neurons as previously reported^47,48^. Examining all cells, we observed a slight but significant decrease in SFARI gene scores within *Adnp* cKOs. Gene scores were not significantly affected in cHets (Fig. 8F, G). However, when we examined upper-layer neurons specifically, SFARI gene scores were significantly reduced in both *Adnp* cHets and cKOs, as well as in *Chd4* cKOs (Fig. 8F, H). We next examined the expression of SysNDD genes. We noted that SysNDD genes were also upregulated in neurons, albeit to a lesser extent (Fig. 8I, J). We observed significant reductions in SysNDD gene scores in *Adnp* and *Chd4* cKOs across all cells, as well as in upper-layer neurons (Fig. 8K). The co-regulation of NDD gene networks by Adnp and Chd4 suggests a mechanism that could potentially contribute to the etiology of ASD/ID.

## Discussion

To characterize functions for the ChAHP complex, previous investigations have focused on *ADNP* mutant models based on the argument that ADNP is an exclusive and essential component of ChAHP^2,15^. However, *ADNP* mutants arrest at the onset of neurodevelopment^2,20,29,30,49,50^, precluding genetic analysis of later processes. To bypass early lethality, we generated telencephalon-specific *Adnp* and *Chd4* cKOs. We found that these were grossly indistinguishable, albeit that *Chd4* cKOs were not obtained in Mendelian ratios^26^. Co-immunoprecipitations revealed that Adnp and Chd4 proteins readily associated *in vivo*. Genomic and transcriptomic profiling suggested that Adnp and Chd4 act cooperatively to regulate the expression of a subset of genes involved in cortical growth. Thus, while ADNP has been linked to a variety of molecular mechanisms^3^, our data indicate that Adnp acts via the ChAHP complex to control neocortical progenitor proliferation. Our study thus reveals a role for the ChAHP complex in the production of upper-layer cortical neurons, and sheds new light on the linkage between chromatin remodelling and NDDs.

### Adnp licenses apical progenitor proliferation to expand upper neocortical layers

Cortical neurogenesis is clearly linked to NDD etiology – especially with respect to ID, where microcephaly and macrocephaly are common comorbidities. The specific requirement for Adnp in upper-layer neuron production was nonetheless surprising given Adnp’s generally ubiquitous expression pattern. *Adnp* transcript was previously shown to be elevated in upper cortical layers^35^, which we confirmed using scRNA-seq and immunohistochemistry. Nonetheless, the defect in cortical neuron production was instead traced to neocortical progenitors and proved to be context-dependent. *Adnp* cKOs exhibited proliferation defects in apical rather basal progenitors, and at late but not early stages. However, at the transcriptional level, gene expression imbalances were surprisingly consistent across timepoints and cell types, suggesting that inherent differences in cellular vulnerability might explain the context-specific phenotypes. For example, progenitor hypoproliferation might arise during late stages due to cumulative defects that build up progressively over time, while apical progenitors might be specifically affected because they self-renew much more extensively versus basal progenitors^32^.

Interestingly, several recent studies have linked NDD etiology to defects in the genetic programs that establish neocortical layers. For example, key transcription factors that determine neuronal subtype identities in the neocortex have been genetically associated with ASD^51^. Moreover, transcriptomic profiling has revealed that as a group, ASD risk genes are particularly well expressed in upper-layer cortical neurons^47,48^. Perhaps for this reason, upper-layer cortical neurons were shown to be particularly vulnerable to perturbations in ASD risk genes^52^. ID risk genes are also highly expressed in excitatory neurons, but are additionally well expressed in proliferating progenitors during mid-gestation^53^. ID risk genes might therefore act at slightly earlier steps during neurogenesis, resulting in more severe phenotypes. Consistent with this model, our data map ChAHP functions to cortical progenitors – consistent with the genetic association between *ADNP*, *CHD4*, and ID. However, our scRNA-seq data additionally show that in neurons, ChAHP has an important role in regulating a subset of NDD risk genes, which might explain why *ADNP* occupies such a prominent position in the genetic landscape of ASD.

Like Adnp, several other broadly expressed chromatin remodellers such as Arid1a, Atrx, Atxn1, and Cic have been shown to be required for the production or maintenance of particular neocortical layers^54–57^. The selective impact of *Adnp* deletion on upper-layer neuron production also resembles *Pou3f2* and *Pou3f3* (*Brn-1* and *Brn-2*) double mutants, which exhibit major deficits in progenitor proliferation beginning at E14.5^58^. Accordingly, we found that the expression *Pou3f1/2/3* transcripts and other upper-layer marker genes was systematically reduced in *Adnp* and *Chd4* cKOs. However, reductions in Pou3f2 levels were restricted to cortical plate neurons, and were not apparent in progenitors. Moreover, *Pou3f2/3* double mutants were reported to exhibit defects in intermediate progenitor proliferation^58^, whereas proliferation defects mapped to apical progenitors in the *Adnp* cKO. The downregulation of *Pou3f* factors is therefore unlikely to completely explain the *Adnp* cKO phenotype. Indeed, we found that Adnp and Chd4 cooperatively regulated several additional genes linked to progenitor proliferation, including *Cdk4* and *Ctnnb1*. Across evolution, along with increases in overall brain size and surface area, the evolutionary expansion of the upper cortical layers in humans has been proposed as a key substrate for the emergence of higher-order cognitive functions^59,60^. While the mechanisms that scale neocortical layers are not well understood, our data suggest that the ChAHP complex may participate in this process.

Although the neocortex is only found in mammals, *Adnp* functions in brain development have previously been examined in other model organisms. In zebrafish, *adnpa/adnpb* double mutants or *adnpa/adnpb*; *adnp2a/adnp2b* morphants were viable and the neural tube was established correctly^20,50^. Mutants or morphants exhibited apparent reductions in brain size and/or neural marker gene expression that were attributed to misregulation of Wnt signaling and consequent apoptosis. Stabilizing interactions between beta catenin and Adnp were demonstrated, along with extensive relocalization of Adnp protein into the cytoplasm of neural progenitors and neurons. However, we think that beta catenin stabilization is unlikely to underlie the cortical growth defects observed in the *Adnp* cKO. While beta catenin drives proliferation, it is additionally linked to the specification of lower-layer neurons in the neocortex^61–63^. By contrast, we find that Adnp is required for upper-layer neuron production. Moreover, unlike the zebrafish ectoderm^20^, we did not observe cytoplasmic localization of Adnp protein. On the other hand, we did observe that Adnp and Chd4 were co-recruited to the promoter of the beta catenin gene itself (*Ctnnb1*), suggesting that Adnp might regulate beta catenin levels transcriptionally rather than post-translationally. These discrepancies might reflect contextual differences between Adnp functions in the early ectoderm versus neural progenitors, or perhaps evolutionary differences between zebrafish and mice. On the other hand, in *Xenopus* embryos, *Adnp* knock-down via CRISPRi led to a marked reduction in the proliferation of telencephalic progenitors^64^, which corresponds very well with the neocortical phenotype that we observe - albeit that the murine phenotype was restricted to late stages of neurogenesis.

The stage-specific defects in neuronal production observed in *Adnp* cKOs, combined with the precocious upregulation of gliogenic genes might suggest a linkage between ChAHP and epigenetic developmental timing mechanisms. Indeed, other chromatin remodelling complexes have been linked to progenitor competence transitions, including the NuRD and polycomb complexes, as well as DNA methylation^65–68^. Crosstalk between these complexes has been well-documented^68–70^, and will be important to address in future work.

### The ChAHP complex and neurodevelopmental gene regulation

Adnp has been linked to heterochromatin formation and gene repression in some contexts^2,71^. However, in the brain, biochemical and genomic profiling revealed that Adnp instead localizes to euchromatic elements, where it associates with Cbx3 (HP1γ) and Pogz^18^. Like *ADNP*, *POGZ* is also among the most commonly *de novo* mutated genes associated with ASD and ID^4,7,8,72–74^. In mice, *Pogz* mutants exhibit relatively subtle deficits in cortical growth^18,75^, but upper-layer neuron production was shown to be affected in one study^76^. Adnp has additionally been shown to occupy active genes where it resolves R-loops, which are RNA/DNA triplex structures that can interfere with transcription^30^. Our data are consistent with a euchromatic role for Adnp – perhaps at R-loops. Adnp and Chd4 were found to co-occupy active regulatory elements, and most DEGs were downregulated in *Adnp* cKOs. However, the strong phenotypic and molecular resemblance between *Adnp* and *Chd4* cKOs argues that with respect to cortical development, functionally important interactions between Adnp and R-loops, or other protein partners like Pogz - likely occur within the context of ChAHP. For example, Pogz could conceivably function as an accessory subunit of ChAHP. Perhaps Chd4 recruitment might augment Adnp’s ability to resolve R-loops^30^. Indeed, other chromatin remodellers have been shown to regulate R-loops, including Atrx and BAF^77,78^. Previous biochemical and immunohistological profiling revealed that Chd4 is critical for the assembly of the NuRD complex in cortical progenitors, since Chd3/5 are mainly expressed in differentiating neurons^13^. The close phenotypic resemblance exhibited by murine *Adnp* and *Chd4* cKO brains was therefore surprising, and suggested that Chd4 regulates progenitor proliferation via ChAHP rather than NuRD. On the other hand, *Adnp* and *Chd4* cKOs were not identical. *Chd4* cKOs were not recovered in Mendelian ratios and additionally exhibited sporadic hydrocephalus^26^. Moreover, transcriptomic data revealed that Adnp and Chd4 regulated a subset of genes oppositely, including the *Chd4* paralogs *Chd3* and *Chd5*, as well as immediate early genes like *Fos* and *Jun*. Interestingly, many of these genes were previously shown to be NuRD targets in the cerebellum^28^. One possible explanation is that Adnp might sequester Chd4 within ChAHP, and thereby limit Chd4’s availability for incorporation into the NuRD complex. However, in the *Adnp* cKO, cut&run-seq data did not reveal increased recruitment of Chd4 to target loci, and co-immunoprecipitations did not reveal increased association between Chd4 and other NuRD subunits (data not shown). We also found that many oppositely regulated genes were directly occupied by both Adnp and Chd4 rather than by Chd4 alone. It may be that both ChAHP and NuRD regulate these loci oppositely. Future work will be required to decipher the precise mechanisms responsible for these effects.

While *Adnp* and *Chd4* cKO models can provide insight into regulatory functions that could in turn be related to ASD/ID etiology, there are important caveats to our study. First, the strong cKOs generated for this study may not be a good model for the *de novo* heterozygous mutations linked to human NDDs. Indeed, we recently showed that *Chd4* cHets and cKOs are phenotypically divergent from constitutive *Chd4* heterozygotes that more closely match human NDD mutations^26^. Indeed, only 20 genes were found to be differentially expressed in *Adnp* cHets versus wild-type, although a few DEGs were potentially interesting, including the immediate early genes *Fos* and *Jun*. Additional DEGs included *Nfix*, *Kif5c*, and *Phip*, which are orthologous to SFARI ASD risk genes. An additional limitation is that we focused our genomic and transcriptomic profiling on embryonic stages of development. Indeed, a recent study performed transcriptomic profiling of brains from germline *Adnp* heterozygotes at juvenile and adult stages^49^. Few if any DEGs appear to be common between the two studies. It may be that Adnp genomic occupancy is drastically remodelled during postnatal development. Despite these limitations, the characterization of ChAHP functions during neocortical development should provide a strong foundation for deciphering how pathogenic mutations affect neurodevelopment, and thereby underlie NDD etiology.

## Methods

### Animals

Animal work was performed in accordance with the guidelines of the Canadian Council on Animal Care and the uOttawa Animal Care and Veterinary Service, using approved ethical protocols OHRI-2856; OHRI-2867; OHRI-3155; and OHRI-3949. The *Adnp^Flox^* allele was generated by Biocytogen (Wakefield, MA) and confirmed using long-range PCR and Southern blotting. Oligos for CRISPR, Southern probes, and genotyping are listed in Table S2. During embryonic stages, animals of either sex were used interchangeably (see also Fig. S5). Postnatal animals were analyzed according to sex separately, and pooled post-hoc. Both *Adnp^Flox^* and *Chd4^Flox^* animals were backcrossed onto the C67BL6/J background.

### Brain Measurements

Dissected brains were imaged using a Zeiss Stemi 508 stereo microscope with Axiocam ERc 5s camera. Images were imported into Fiji (ImageJ), and the outline of the dorsal telencephalon was traced using the “Freehand selections” tool. Brain weights were measured using an XS105 analytical balance (Mettler Toledo).

### Histology and microscopy

Immunohistochemistry was performed as previously described^37^, except that antigen retrieval was omitted. Briefly, freshly dissected brains were fixed via immersion in 4% paraformaldehyde/PBS. Tissues were cryoprotected overnight in 20% sucrose (w/v)/PBS, followed by a 1:1 mixture of 20% sucrose/PBS and OCT Tissue-Plus™ O.C.T. Compound (23-730-571; Fisher Scientific) overnight. Brains were embedded and flash-frozen in 1:1 sucrose/O.C.T., and then cryosectioned at 12 microns (embryonic stages) or 16 microns (postnatal stages). Primary antibodies are listed in Table S3. Confocal microscopy was performed using a Zeiss LSM900 instrument (CBIA Core, uOttawa) using Zen (Zeiss), Fiji (ImageJ), and Adobe Photoshop (Adobe) software. EdU staining was performed using the Click-iT™ EdU Cell Proliferation Kit (Invitrogen). All cell counts were performed manually and unblinded in the rostral somatosensory cortex. All images were processed using linear transformations, except the images in Fig. 3A-C and 4A-C.

### Immunoprecipitation and western blot

P3 neocortex or hippocampus was freshly microdissected and sonicated in RIPA buffer (10 mM Tris pH 8.0; 140 mM NaCl; 1mM EDTA; 0.5 mM EGTA; 1% Triton X-100; 0.1 % deoxycholate; 0.1 % SDS) supplemented with cOmplete™ EDTA-free Protease Inhibitors (11836170001; Roche). 30 µl of protein A or G paramagnetic beads (73778S or 9006S; New England Biolabs) were washed and then complexed with primary antibodies or naïve rabbit IgG at 4°C with end-over-end rotation. Beads were washed and 2 mg of protein lysate was added to each tube. Lysates were diluted to 1 ml with RIPA buffer with protease inhibitors and incubated overnight at 4°C with end-over-end rotation. Next, beads were pipetted into fresh tubes, and then washed 3x 30 min with RIPA buffer. Proteins were eluted by incubating the paramagnetic beads with Laemmli Sample buffer (1610747; Bio-Rad) supplemented with 2-mercaptoethanol, and boiling for 10 minutes.

Proteins were separated on SDS-PAGE gels and transferred to PVDF membranes (IPVH00010; Fisher Biosciences) using a Trans-Blot® Turbo™ Transfer System (Bio-Rad). After transfer, membranes were post-fixed in 95% ethanol. Membranes were then transferred into TBST (100mM Tris pH 8.0, 150mM NaCl, 0.5% Tween 20), and then blocked in Blotto (TBST supplemented with 5% w/v skim milk powder). Primary antibodies (see Table S3) were diluted in Blotto and applied to the membranes overnight at 4°C with end-over-end rotation. Membranes were then washed in TBST. HRP-conjugated secondary antibodies (see Table S3) were applied for 1 h in Blotto. After washing, westerns were visualized with the Clarity™ Western ECL Substrate (Bio-Rad) and a Chemidoc MP system (Bio-Rad).

### Cut&run-seq

Cut&run-seq was performed using the Cutana cut&run kit (14-0500; Epicypher), following the manufacturer’s instructions. Briefly, embryos were harvested and the dorsal telencephalon was microdissected. Cells were dissociated with mechanical trituration in order to retain the glycoproteins necessary for adhering to the concavanalin beads. 500 000 cells were used in each reaction. Cut&run antibodies are listed in Table S3. Libraries were prepared using the NEBNext® Ultra^TM^ II Library Prep Kit (New England Biolabs). Paired-end 150 sequencing was performed using the NextSeq 500 platform (Illumina) to a read-depth of ∼20-35 million reads per sample.

Bioinformatic analysis was performed using Galaxy^79^. FASTQ files were processed via Fastq Groomer^80^ and Trimmomatic^81^, and then mapped to the mm9 genome using Bowtie2^82^. To normalize samples to the *E. coli* spike-in, the trimmed reads were mapped to the *E. coli* K12 genome using Bowtie2. Coverage was calculated using Samtools Flagstat. Mapped BAM files were then scaled using Deeptools^83^. See Table S4 for scaling information. Peak calling was performed with Macs2^84^, or SEACR^85^, and we used GREAT^86^ for peak-to-gene annotation. Adnp and IgG cut&run-seq data previously generated by the John Rubinstein lab^18^ were obtained from <u>Series GSE187010</u> and mapped *de novo* as per above to obtain called peaks. H3K4me3 (https://www.encodeproject.org/files/ENCFF053SSJ/) and H3K27ac (https://www.encodeproject.org/files/ENCFF714RJJ/) ChIP-seq data generated by the Bing Ren lab^43^ were obtained from the ENCODE Consortium^87^. Genome-wide histograms were generated using Seqplots^88^. Overlapping peaks were identified using Intervene^89^. Cut&run-seq data will be deposited in the GEO database.

### Multi-seq

Embryos were harvested and the dorsal telencephalon was microdissected. Cells were dissociated with papain (Worthington) in the presence of DNAse I, washed, and then barcoded. Briefly, 250 000 dissociated cells per replicate were incubated with ‘anchor’ and ‘co-anchor’ lipid-modified oligonucleotides generously provided by the Gartner lab^39^. Each replicate was then co-incubated with barcode oligonucleotides for 10 minutes and then washed 3 times with PBS. Replicates were then pooled in equivalent ratios, and approximately 20 000 pooled cells were combined in individual Chromium™ runs (3’ Library & Gel Bead Kit v2, PN-120237, 10X Genomics). Expression library FASTQs were processed using CellRanger (10X Genomics). The deMULTIplex workflow^39^ was used to remove doublets and exclude cells lacking barcodes.

Output files were filtered and analyzed using Scanpy version 1.9.1^90^ in Python (Python Core Team n.d.). Genes detected in less than 3 cells were removed from the analysis. Contaminant or low-quality cells (less than 5000 genes detected, less than 20 000 reads/counts detected or more than 0.05% of mitochondrial genes detected) were also excluded. Differential gene expression analyses were performed using or Scanpy (Wilcoxon signed-rank test), or MAST version 1.24.0^42^. In order to avoid batch effects, pairwise comparisons were made between cKO samples and littermate control samples that had been processed together in the same batch. Littermate *Chd4* cHet samples were used as paired controls for E16.5 *Chd4* cKOs, and littermate *Adnp* cHet samples were compared with E18.5 *Adnp* cKOs. GO terms analysis was performed using ShinyGO^91^. Gene scoring was generated using Scanpy (scanpy.tl.score_genes). Ensembl BioMart^92^ was first used to obtain mouse orthologues of human SFARI genes^45^, or “definitive” SysNDD genes^46^. Orthologues could not be obtained for 7 SFARI genes and 14 SysNDD genes. Statistical analysis of the gene score results was performed using Scipy.stats. scRNA-seq data will be deposited in the GEO database.

### Quantitation and statistical analysis

For cell count data and brain measurements, statistical analyses were performed using GraphPad Prism software. All tests were 2-tailed. n-values refer to biological replicates (different animals). All bar graphs display meanLJ±LJSEM. Sample sizes were not predetermined by power calculations. Venn diagrams were generated using the VennDiagram Package in R. ANOVAs passed the Shapiro-Wilk test for normality and the Brown-Forsythe test for variance. All error bars are mean ± SEM. We did not exclude any datapoints in this study, with the exception of cells that did not meet quality control standards in our scRNA-seq analyses.

## Supporting information

Supplemental Section

## Acknowledgements

We thank Ryan MacDonald for comments on a previous version of this manuscript. We thank Katia Georgopoulos and Toshimi Yoshida for sharing the *Chd4^Flox^* mice. We thank Ivana Herrera, Whitney Carter, and Ida Malloy for help in initiating this project. We thank Zev Gartner, Chris McGinnis, David Cook, and Barbara Vanderhyden for generously providing Multi-seq protocols, reagents, and advice. We thank the laboratories of Bing Ren and John Rubinstein for providing genomic datasets upon which this study relied. For scRNA-seq and cut&run-seq experiments, we also thank Katayoun Sheikheleslamy, Caroline Vergette, and Pearl Campbell from the *Stemcore Molecular Biology Core facility*, as well as Chris Porter from the *OHRI Bioinformatics Core Facility*. We thank Chloë van Oostende-Triplet and the *Cell Biology and Image Acquisition Core Facility*, the staff of the uOttawa *Animal Care and Veterinary Service*.

We thank members of the Mattar and Picketts labs for their ongoing help and input. This work was generously supported by a SFARI Pilot Award (609562 to PM) as well as a Young Investigator Award from the Brain & Behavior Research Foundation (28564 to PM). We also acknowledge generous support from the Canadian Institutes of Health Research (CIHR) Operating Grants (PJT-166032 (PM), PJT-166074 (PM) PJT-159619 (DJP), PJT-165994 (DJP), and PJT-180577 (DJP)), as well as from the Society of Hellman Fellows (to KS) and McKnight Foundation (to KS). We additionally acknowledge the generous support of the New Frontiers in Research Fund (NFRFT-2022-00327 to PM). The project was also supported by an infrastructure grant from the Canada Foundation for Innovation for confocal microscopy (JELF 37688 to PM). PM gratefully holds the Gladys and Lorna J. Wood Chair for Research in Vision.

## Author Contributions

Study design: PM; Investigation: SC-D, SL, SM, RS, ZEH, SVJ, and PM; Bioinformatic analysis; XS, JALF, and PM; Data analysis; SL, XS, JALF, and PM; Supervision and funding acquisition: DJP, KS, and PM; Writing—original draft: PM; Writing, review, and editing: all authors.

## Availability of Data and Materials

Cut&run-seq and scRNA-seq data will be uploaded to the GEO repository. *Adnp^Flox^* mice will be deposited with the Jackson Laboratories for distribution. Microscopy datasets are available from the corresponding author on request.

