## Supplemental Section for "The ChAHP chromatin remodelling complex regulates neurodevelopmental disorder risk genes to scale the production of neocortical layers"

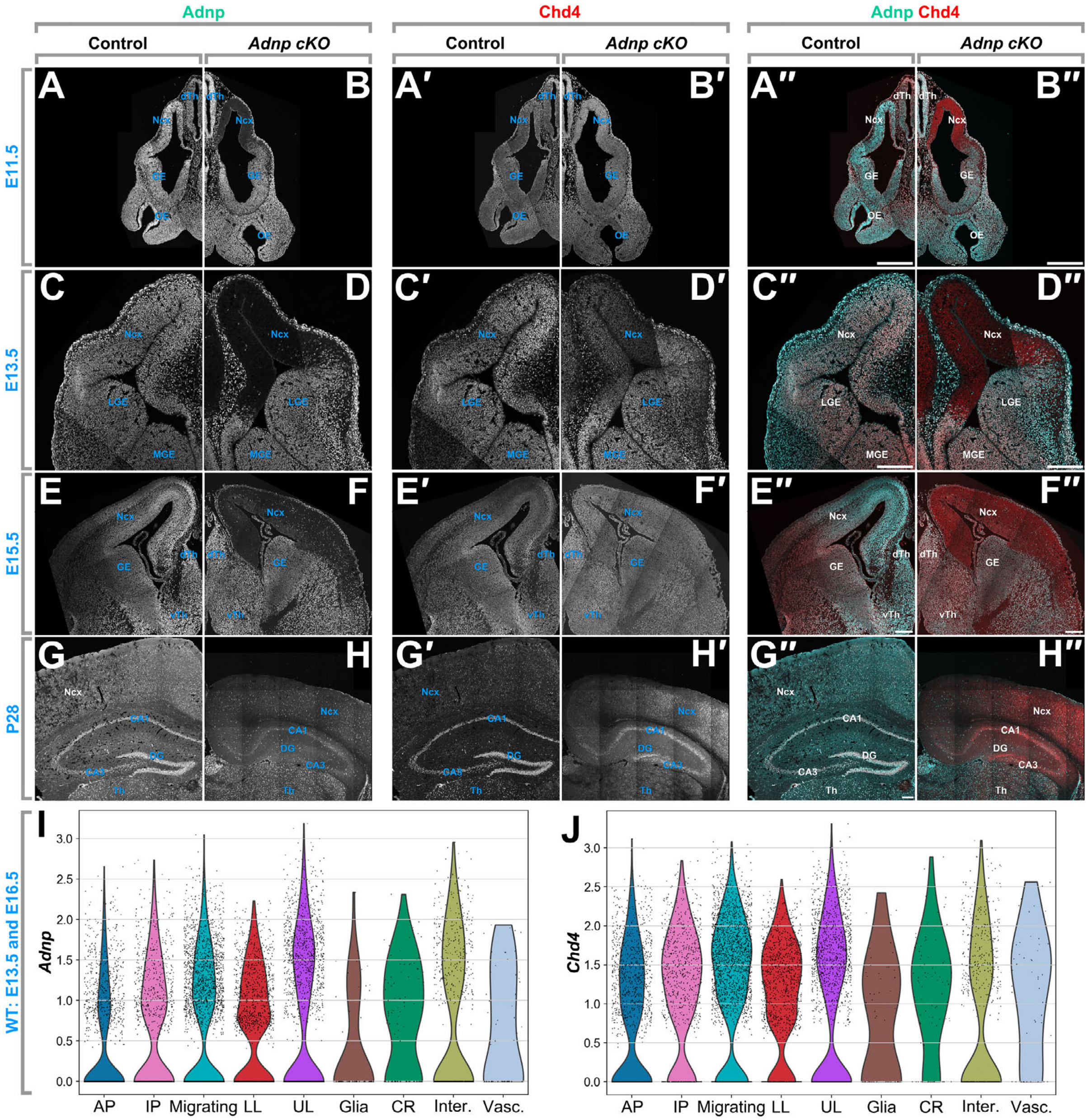


Figure S1. Adnp and Chd4 are broadly co-expressed during neurodevelopment. (A-H) Co-immunohistochemistry for Adnp (A-H), Chd4 (A′ -H′), or merged (A″-H″) from control (A, C, E, G) or *Adnp* cKO brains (B, D, F, H). Stages are as indicated. (I, J) scRNA-seq expression data for annotated cell types from E13.5 and E16.5 wild-type samples. *Adnp* (I) or *Chd4* (J). Ncx: neocortex; GE: ganglion eminence; LGE: lateral ganglionic eminence; MGE: medial ganglionic eminence; Th: thalamus; dTh: dorsal thalamus; vTh: ventral thalamus; OE: olfactory epithelium; DG: dentate gyrus; CA1: Cornu Ammonis 1; CA3: Cornu Ammonis 3. AP: apical progenitors; IP: intermediate progenitors; LL: lower-layer neurons; UL: upper-layer neurons; CR: Cajal-Retzius cells; Inter.: interneurons; Vasc.: vascular endothelial cells. Scale bars = 200 µm.

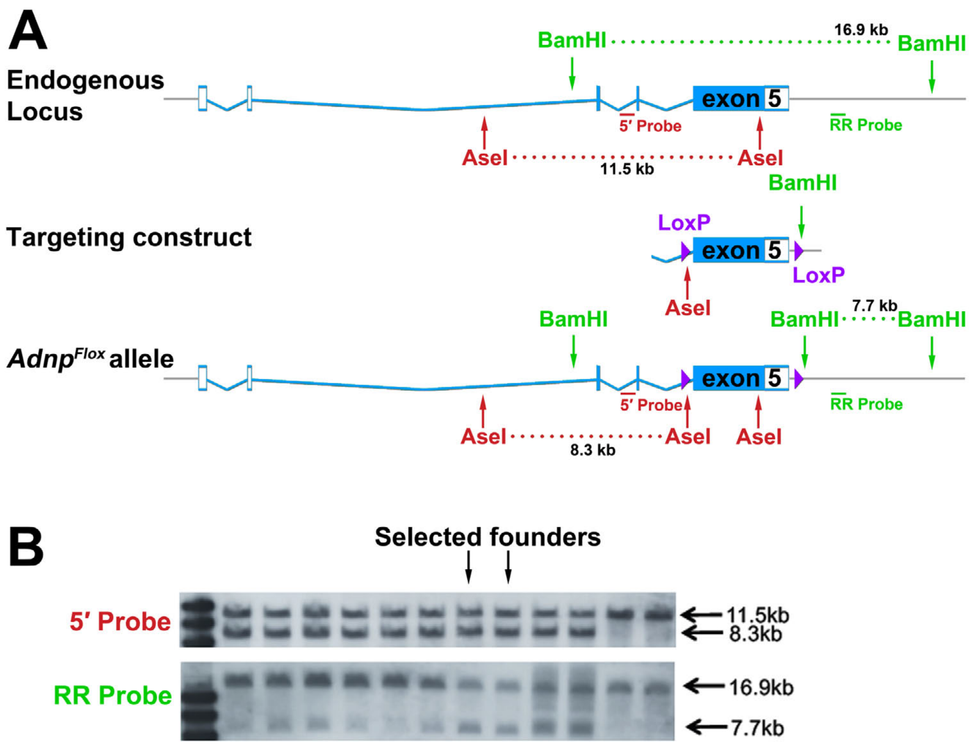


Figure S2. Generation of the *Adnp*$^{Flox}$ allele. (A) Schematic of CRISPR-HDR strategy. PAM sites were selected at the ends of the targeting construct. Red and green arrows/text show the position of indicated restriction enzyme sites and probes utilized for Southern blotting. (B) Southern blot analysis of F1 mice generated via the CRISPR strategy. The indicated founder animals were selected for subsequent breeding.

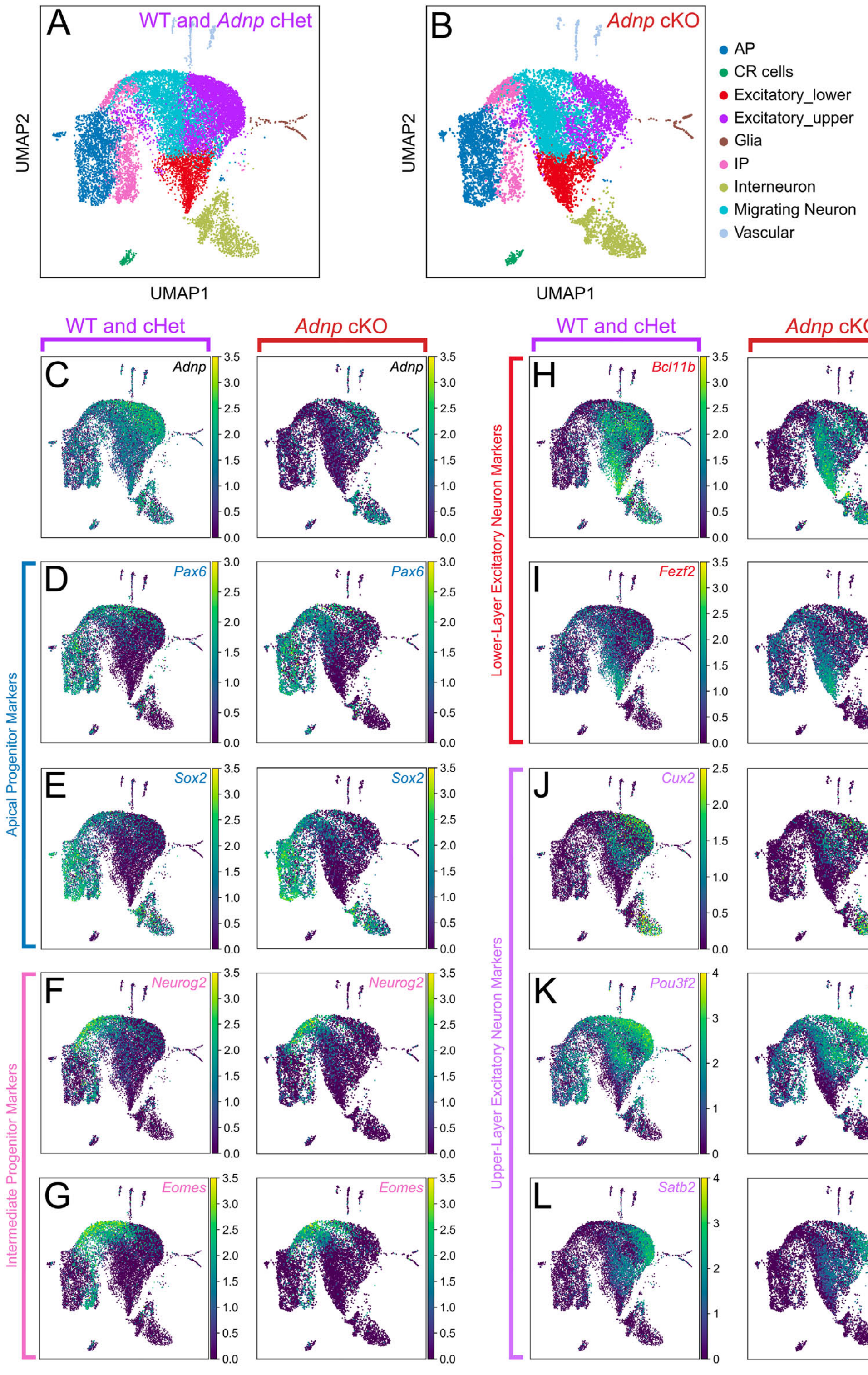


Figure S3. Comparison of marker gene expression in *Adnp* cKOs versus controls. (A) Cell type annotation of WT and *Adnp* cHet cells replicates. (B) Cell type annotation of *Adnp* cKO replicates. (C) Expression of *Adnp* in WT and *Adnp* cHet replicates (left) or *Adnp* cKO replicates (right). (D-L) Marker gene expression of *Adnp* in WT and *Adnp* cHet replicates (left) or *Adnp* cKO replicates (right). (D, E) Apical progenitor markers *Pax6* (D) and *Sox2* (E). (F, G) Intermediate Progenitor markers *Neurog2* (F) and *Eomes* (G). (H, I) Lower-Layer neuron markers *Bcl11b* (H) and *Fezf2* (I). (J-L) Upper-layer neuron markers *Cux2* (J), *Pou3f2* (K), and *Satb2* (L).

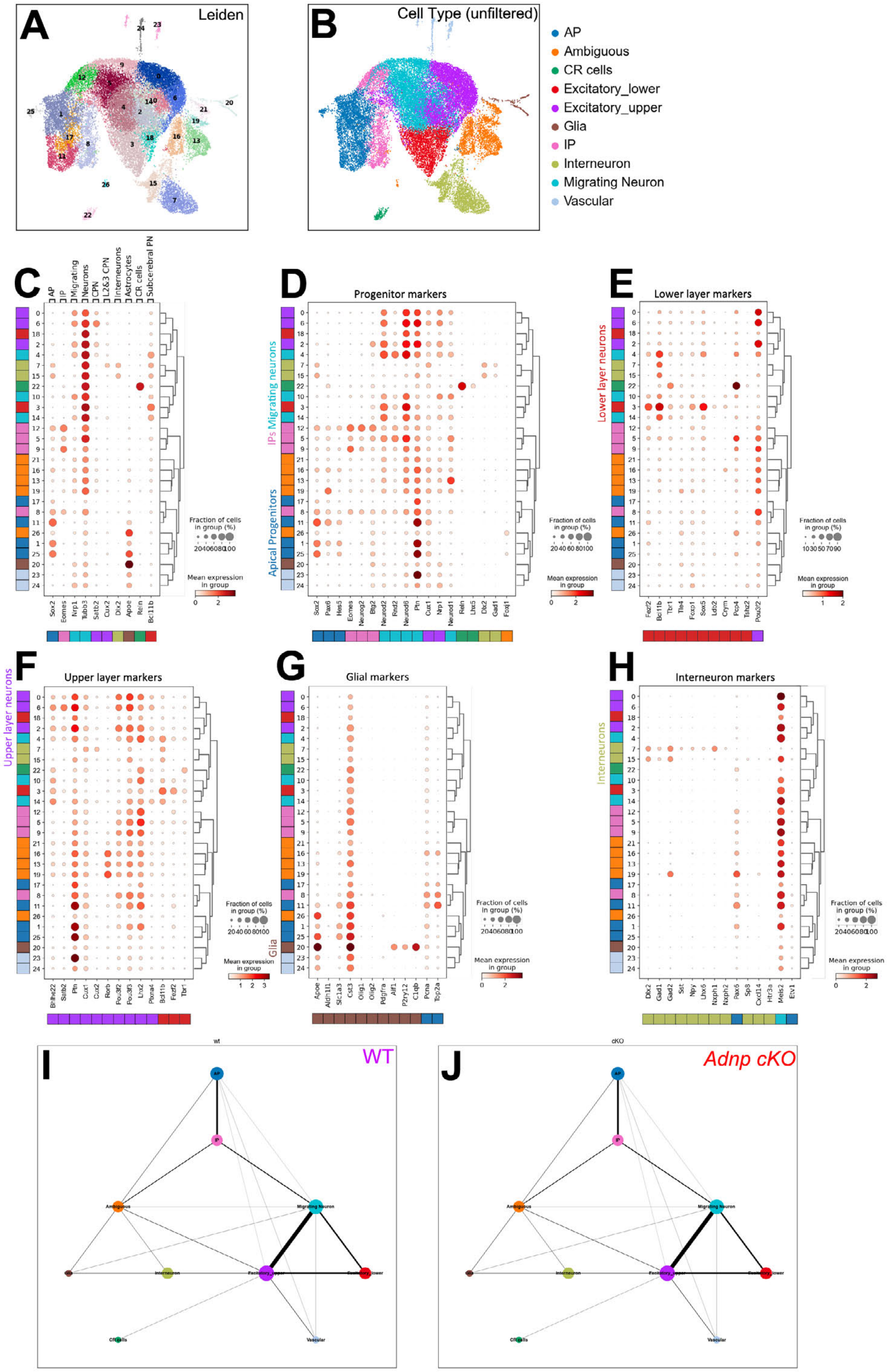


Figure S4. Cell-type annotation. (A) Cluster identification via the Leiden algorithm. (B) Assigned cell types. Annotation was performed using a previously published gene expression framework[1]. (C-H) Dotplots of marker gene expression for Leiden clusters. Color bars under the X-axes refer to the cell-type-specific expression pattern of the corresponding gene. Color bars on the Y-axes refer to the assigned cell category for each Leiden cluster. (I, J) Trajectory inference for wild-type (I) or *Adnp* cKO (J) using PAGA[3], with apical progenitors set as the starting point. Line thickness indicates the strength of the inferred trajectory. AP: apical progenitors; IP: intermediate progenitors; CR: Cajal-Retzius cells.

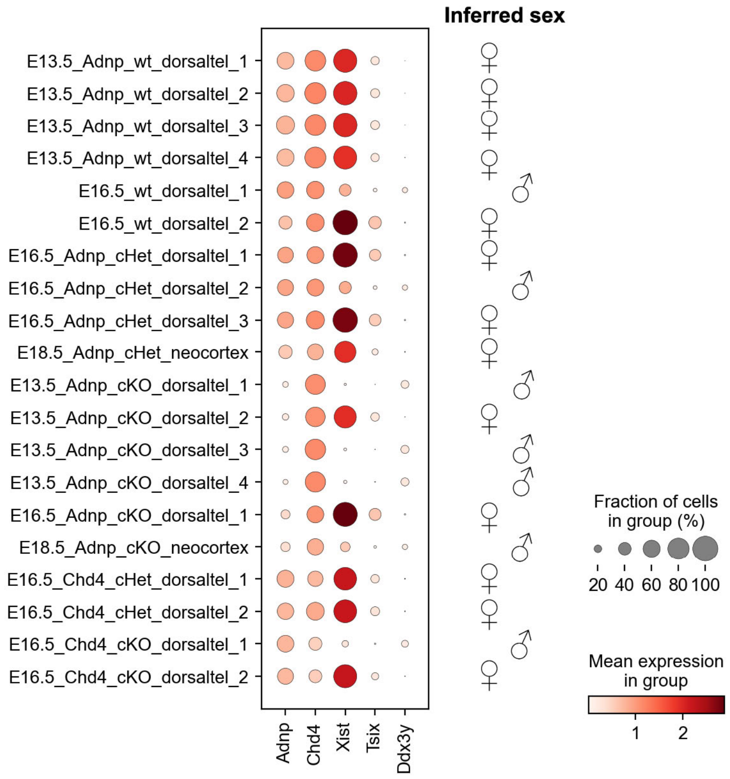


Figure S5. Analysis of gene expression in individual scRNA-seq replicates. Dotplots of gene expression grouped by genotype. The expression of *Adnp* and *Chd4* transcripts validates the conditional knockout samples. The expression of *Xist* and *Tsix* transcripts from the inactivated X chromosome is elevated in female samples, while the expression of *Ddx3y* from the Y chromosome is male-specific.

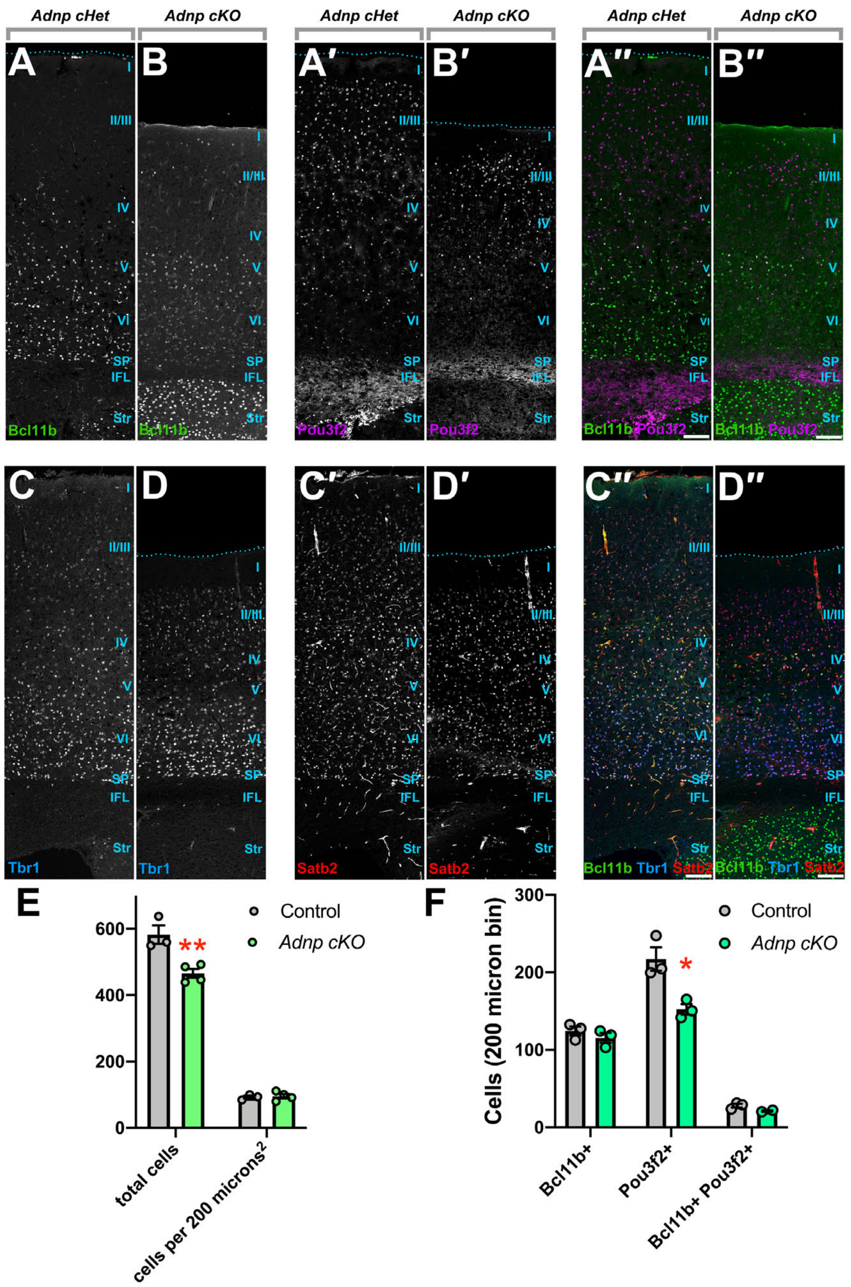


Figure S6. *Adnp* is required for the expansion of upper-layer neocortical neurons. (A, B) Expression of the lower-layer marker Bcl11b and the upper-layer marker Pou3f2 at P28. (C, D) Expression of Tbr1 and Satb2 at P28. (E) Quantitation of total neocortical cells in a 200 μm wide bin, and cell density per 200 $\mu m^2$, counted in the rostral somatosensory cortex. ** p = 0.0090 by Student's t-test. (F) Quantitation of total Bcl11b+, Pou3f2+, and double-positive neurons in a 200 μm wide bin. * p = 0.0175 by Student's t-test. Ncx: neocortex; Str: striatum; IFL: inner fibre layer; SP: subplate. Roman numerals indicate neocortical layers. Scale bars = 100 μm.

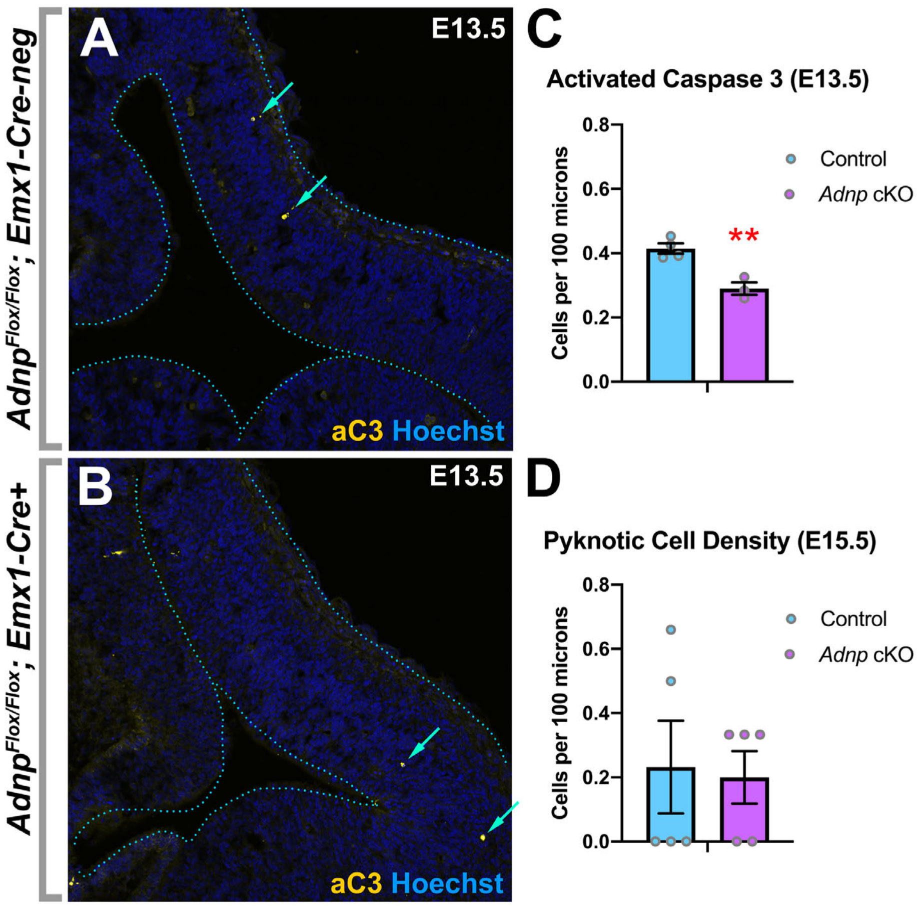


Figure S7. Adnp is not required for cell survival during neurogenesis. (A, B). Immunohistochemistry for activated caspase 3 in E13.5 control (A) or *Adnp* cKO (B) neocortices. Arrows indicate apoptotic cells. (C) Activated caspase 3 counts at E13.5. (D) Counts for pyknotic nuclei at E15.5. ** p = 0.0039 by Student's t-test.

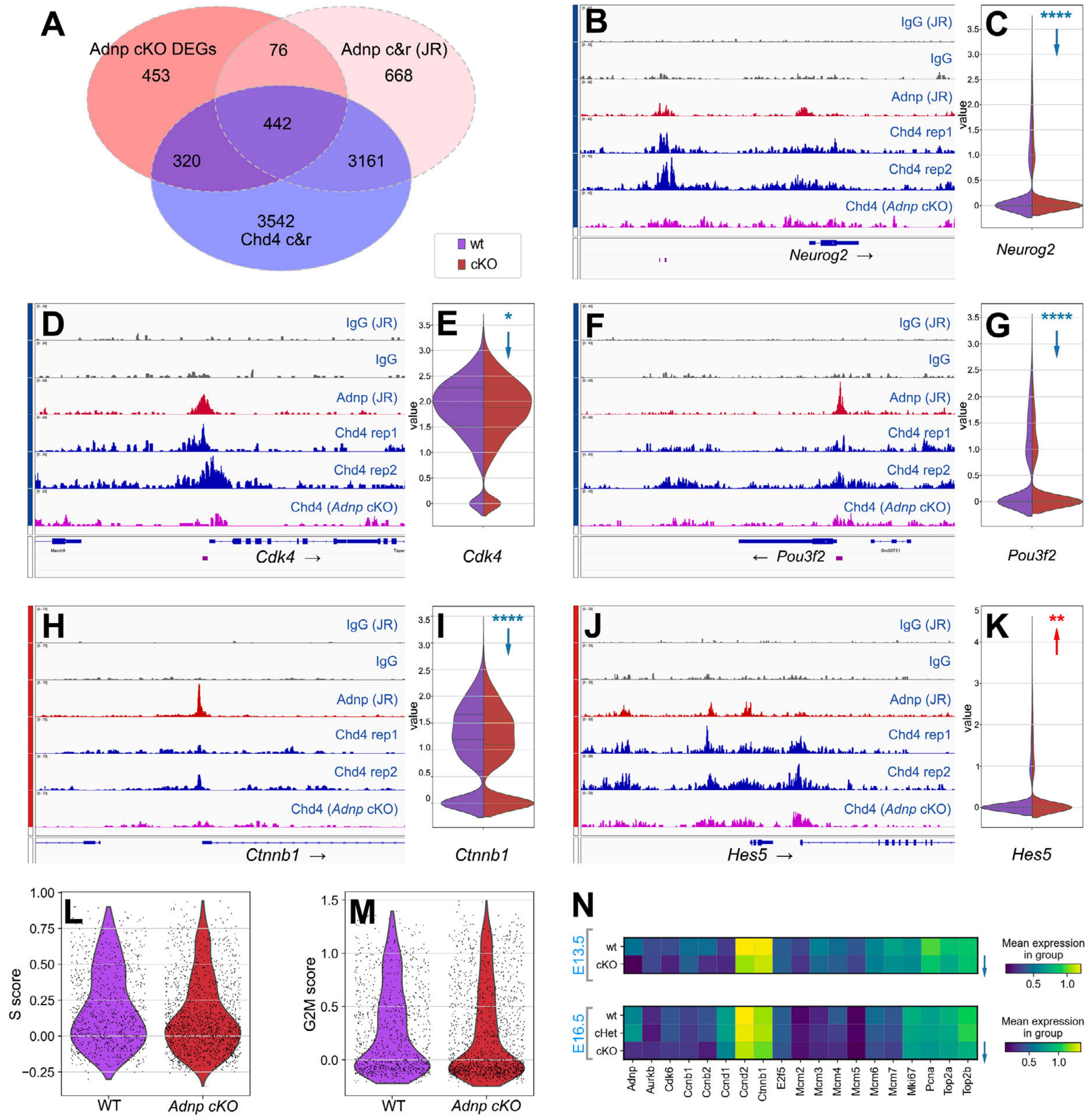


Figure S8. Adnp and Chd4 co-occupy a subset of *Adnp* cKO DEGs. (A) Venn diagram comparing DEGs identified E13.5 *Adnp* cKOs (red) with genes associated with E13.5 Chd4 cut&run-seq peaks (blue) or E13.5 Adnp cut&run-seq peaks (pink). (B-K) Genomic views of selected genes co-occupied by Adnp and Chd4 (see A), with violin plots of mRNA expression from E13.5 WT (purple) or *Adnp* cKO (red) apical progenitors. All cut&run-seq tracks are group-autoscaled. Adnp cut&run-seq and matched IgG control datasets were previously generated by the John Rubinstein lab (JR)[2]. (L) S-phase or (M) G2/M-phase cell cycle scores from E13.5 apical progenitors as indicated. (N) Matrix plots of selected proliferation genes from E13.5 or E16.5 apical progenitors as indicated. * $p < 0.05$, ** $p < 0.01$, *** $p < 0.001$, **** $p < 0.0001$ by adjusted p-value (Wilcoxon rank sum test).

**Table S1:** scRNA-seq metrics

| Stage | Genotype | Sample ID | Cell number (post QC) | Batch | Mutli-seq Barcode |
|---|---|---|---|---|---|
| E13.5 | (4 wt; 4 *Adnp* cKO) | all E13.5 | 12208 | 1 | |
| E16.5 | (2 wt; 3 *Adnp* cHet; 1 *Adnp* cKO) | (batch 2) | 12157 | 2 | |
| E16.5 | (2 *Chd4* cHet; 2 *Chd4* cKO) | (batch 3) | 3591 | 3 | |
| E16.5 | | all E16.5 | 15748 | 2 and 3 | |
| E18.5 | (1 *Adnp* cHet; 1 *Adnp* cKO) | all E18.5 | 3443 | 3 | |
| E13.5 | wt | E13.5_Adnp_wt_dorsaltel_1 | 1192 | 1 | 9 |
| E13.5 | wt | E13.5_Adnp_wt_dorsaltel_2 | 1358 | 1 | 10 |
| E13.5 | wt | E13.5_Adnp_wt_dorsaltel_3 | 1107 | 1 | 11 |
| E13.5 | wt | E13.5_Adnp_wt_dorsaltel_4 | 1996 | 1 | 12 |
| E13.5 | *Adnp* cKO | E13.5_Adnp_cKO_dorsaltel_1 | 1322 | 1 | 5 |
| E13.5 | *Adnp* cKO | E13.5_Adnp_cKO_dorsaltel_2 | 1626 | 1 | 6 |
| E13.5 | *Adnp* cKO | E13.5_Adnp_cKO_dorsaltel_3 | 1387 | 1 | 7 |
| E13.5 | *Adnp* cKO | E13.5_Adnp_cKO_dorsaltel_4 | 2199 | 1 | 8 |
| E16.5 | wt | E16.5_wt_dorsaltel_1 | 1992 | 2 | 7 |
| E16.5 | wt | E16.5_wt_dorsaltel_2 | 575 | 2 | 11 |
| E16.5 | *Adnp* cHet | E16.5_Adnp_cHet_dorsaltel_1 | 2109 | 2 | 8 |
| E16.5 | *Adnp* cHet | E16.5_Adnp_cHet_dorsaltel_2 | 1362 | 2 | 9 |
| E16.5 | *Adnp* cHet | E16.5_Adnp_cHet_dorsaltel_3 | 1364 | 2 | 12 |
| E16.5 | *Adnp* cKO | E16.5_Adnp_cKO_dorsaltel_1 | 3033 | 2 | 10 |
| E16.5 | *Chd4* cHet | E16.5_Chd4_cHet_dorsaltel_1 | 217 | 3 | 5 |
| E16.5 | *Chd4* cHet | E16.5_Chd4_cHet_dorsaltel_2 | 261 | 3 | 6 |
| E16.5 | *Chd4* cKO | E16.5_Chd4_cKO_dorsaltel_1 | 1301 | 3 | 7 |
| E16.5 | *Chd4* cKO | E16.5_Chd4_cKO_dorsaltel_2 | 1772 | 3 | 8 |
| E18.5 | *Adnp* cHet | E18.5_Adnp_cHet_neocortex | 2185 | 3 | 1 |
| E18.5 | *Adnp* cKO | E18.5_Adnp_cKO_neocortex | 1243 | 3 | 2 |

**Table S2:** Oligonucleotide sequences. Multi-seq barcodes are highlighted in green.

| Name | Purpose | Sequence |
|---|---|---|
| sgRNA-2 | CRISPR | 5′-GGGGGAACATTGGTCT-3′ |
| sgRNA-16 | CRISPR | 5′-TCCAGGGACCAGAGGTG-3′ |
| A-A-F | targeting | 5′-atcgGTCGACAGGGACCAGAGGTGCTGCACAC-3′ |
| A-A-R | targeting | 5′-atcgGTCGACAGGGACCAGAGGTGCTGCACAC-3′ |
| LR-F | targeting | 5′-atcgGAATTCGGAGTGCTCTCCAGAATTGTGTGT-3′ |
| LR-R | targeting | 5′-atcgCTCGAGATTAATGGAACATTGGTCTCTTTCTGTTTCTTC-3′ |
| RR-F | targeting | 5′-atcgACGCGTCCTGACATAGGGGATCCGGAGGGTTTTCTCCATGTCCATAT-3′ |
| RR-R | targeting | 5′-atcgGCGGCCGCGGGTCCCATAACAGTCATTAGGCTG-3′ |
| 5′-Probe-F | Southern | 5′-TGAGGCACAGTCGTCTGAGTGTTTG-3′ |
| 5′-Probe-F | Southern | 5′-TGGTTTCCAGGGTACATCCCTGACT-3′ |
| RR-Probe-F | Southern | 5′-CACTGCCACCTTGCACTTCAGAAAC-3′ |
| RR-Probe-R | Southern | 5′-AGAGGTCAGCATACAGCAGTGAGGA-3′ |
| Adnp-3’F | genotyping | 5′-TGGCGTAACAGTAAAGGAGAAGTAC-3′ |
| Adnp-3’F | genotyping | 5′-AGAGCTTCAGCATGACTTCCAGGTG-3′ |
| Mi-2β+F S | genotyping | 5′-CTCCAAGAAGAAGACGGCAGATCT-3′ |
| Mi-2 INR A | genotyping | 5′-GTCCTTCCAAGAAGAGCAAG-3′ |
| CRE-F | genotyping | 5′-AGGTGTAGAGAAGGCACTTAGC-3′ |
| CRE-R | genotyping | 5′-CTAATCGCCATCTTCCAGCAGG-3′ |
| Multiseq-1 | barcoding | 5′-CCTTGGCACCCGAGAATTCCAGGAGAAGAAAAAAAAAAAAAAAAAAAAAAAAAAAAAA-3′ |
| Multiseq-2 | barcoding | 5′-CCTTGGCACCCGAGAATTCCACCACAATGAAAAAAAAAAAAAAAAAAAAAAAAAAAAAA-3′ |
| Multiseq-3 | barcoding | 5′-CCTTGGCACCCGAGAATTCCATGAGACCTAAAAAAAAAAAAAAAAAAAAAAAAAAAAAA-3′ |
| Multiseq-4 | barcoding | 5′-CCTTGGCACCCGAGAATTCCAGCACACGCAAAAAAAAAAAAAAAAAAAAAAAAAAAAAA-3′ |
| Multiseq-5 | barcoding | 5′-CCTTGGCACCCGAGAATTCCAAGAGAGAGAAAAAAAAAAAAAAAAAAAAAAAAAAAAAA-3′ |
| Multiseq-6 | barcoding | 5′-CCTTGGCACCCGAGAATTCCATCACAGCAAAAAAAAAAAAAAAAAAAAAAAAAAAAAAA-3′ |
| Multiseq-7 | barcoding | 5′-CCTTGGCACCCGAGAATTCCAGAAAAGGGAAAAAAAAAAAAAAAAAAAAAAAAAAAAAA-3′ |
| Multiseq-8 | barcoding | 5′-CCTTGGCACCCGAGAATTCCACTATAGTAAAAAAAAAAAAAAAAAAAAAAAAAAAAAAA-3′ |
| Multiseq-9 | barcoding | 5′-CCTTGGCACCCGAGAATTCCATAAAATCCAAAAAAAAAAAAAAAAAAAAAAAAAAAAAA-3′ |
| Multiseq-10 | barcoding | 5′-CCTTGGCACCCGAGAATTCCAGTATATGTAAAAAAAAAAAAAAAAAAAAAAAAAAAAAA-3′ |
| Multiseq-11 | barcoding | 5′-CCTTGGCACCCGAGAATTCCATAATCAACAAAAAAAAAAAAAAAAAAAAAAAAAAAAAA-3′ |
| Multiseq-12 | barcoding | 5′-CCTTGGCACCCGAGAATTCCAGTAGCACTAAAAAAAAAAAAAAAAAAAAAAAAAAAAAA-3′ |

**Table S3:** Antibodies and dilutions.

| Antigen | Species | Dilution | Supplier | Catalog | Application |
|---|---|---|---|---|---|
| Adnp | Goat | 1:200<br>1:1000 | R&D Systems | AF5919 | IHC,<br>IP/Western |
| Bcl11b (Ctip2) | Rat | 1:200 | Abcam | ab18465 | IHC |
| Chd3 | Rabbit | 1:200 | Fortis | A301-220A-T | IHC |
| Chd4 | Rat | 1:200 | Biolegend | 942302 | IHC |
| Chd4 | Rabbit | 1:1000<br>1:100 | Abcam | ab72418 | IP/Western,<br>cut&run |
| Cleaved Caspase-3 (Asp175) | Rabbit | 1:500 | Cell Signaling | 9579 | IHC |
| Ctcf | Rabbit | 1:100 | Cell Signaling | 3418 | cut&run |
| Eomes (Tbr2) | Rabbit | 1:200 | Abcam | ab23345 | IHC |
| Ki67 | Rabbit | 1:100 | Millipore Sigma | SAB5500134 | IHC |
| Olig2 | Rabbit | 1:500 | Novus | NBP1-28667 | IHC |
| Pax6 | Rabbit | 1:500 | Novus | NBP2-19711 | IHC |
| Pax6 | Rabbit | 1:500 | Proteintech | 12323-1-AP | IHC |
| Pou3f2 (Brn2) | Rabbit | 1:500 | Cell Signaling | 12137S | IHC |
| Satb2 | Mouse | 1:200 | Abcam | ab51502 | IHC |
| Sox2 | Goat | 1:500 | R&D Systems | AF2018-SP | IHC |
| Tbr1 | Rabbit | 1:500 | Cell Signaling | 49661 | IHC |
| anti-goat Alexa Fluor™ 488 | Donkey | 1:1000 | Jackson | 705-547-003 | 2º (IHC) |
| anti-goat DyLight™ 650 | Donkey | 1:1000 | Novus | NBP1-75604 | 2º (IHC) |
| anti-mouse Alexa Fluor™ 555 | Donkey | 1:1000 | Invitrogen | A-31570 | 2º (IHC) |
| anti-mouse Alexa Fluor™ 647 | Donkey | 1:1000 | Invitrogen | A-31571 | 2º (IHC) |
| anti-rabbit Alexa Fluor™ 488 | Donkey | 1:1000 | Jackson | 111-545-003 | 2º (IHC) |
| anti-rabbit Alexa Fluor™ 647 | Donkey | 1:1000 | Jackson | 711-607-003 | 2º (IHC) |
| anti-rat DyLight™ 550 | Donkey | 1:1000 | Invitrogen | SA510027 | 2º (IHC) |
| anti-Goat HRP | Donkey | 1:10 000 | Invitrogen | A16005 | 2º (Western) |
| anti-Rabbit HRP | Donkey | 1:10 000 | GE Healthcare | NA934 | 2º (Western) |

**Table S4:** Cut&run-seq normalization.

| Sample_ID | Sample_Name | Total reads | *E. coli* paired reads | Percent *E. coli* | Scale factor |
|---|---|---|---|---|---|
| 581930 | IgG Ctl2 | 21117228 | 664251 | 3.15% | 0.840090568 |
| 581931 | Abcam Chd4 Ctl1 | 23376868 | 582381 | 2.49% | 0.958188883 |
| 581932 | Abcam Chd4 Ctl2 | 29603134 | 463685 | 1.57% | 1.203470028 |
| 581934 | Abcam Chd4 CKO6 | 26198838 | 616698 | 2.35% | 0.904869158 |
| 581935 | NEB Ctcf Ctl1 | 31100448 | 789706 | 2.54% | 0.706631329 |
| 581936 | NEB Ctcf Ctl2 | 21788282 | 599799 | 2.75% | 0.930363338 |
| 581937 | NEB Ctcf CKO4 | 23785194 | 665555 | 2.80% | 0.838444606 |
| 581938 | NEB Ctcf CKO6 | 26996200 | 558031 | 2.07% | 1 |

**Supplemental datafile 1:** DEGs from *Adnp* and *Chd4* mutants as determined via Scanpy

Tab 1: DEGs from *Adnp* cKOs vs WT: E13.5, E16.5, and E18.5
Tab 2: DEGs from E16.5 *Chd4* cKOs vs *Chd4* cHets
Tab 3: Gene list intersections: E16.5 *Adnp* cKO DEGs vs. *Chd4* cKO DEGs
Tab 4: DEGs from E16.5 *Adnp* cHets vs WT
Tab 5: Gene list intersections: E16.5 *Adnp* cKO DEGs vs. *Adnp* cHet DEGs
Tab 6: DEGs from annotated interneurons: *Adnp* cKOs vs WT at E13.5

**Supplemental datafile 2:** DEGs from E13.5 *Adnp* cKOs (vs. WT) as determined via MAST[4]

**Supplemental datafile 3:** SFARI and SysNDD gene lists intersected with DEGs from *Adnp* and *Chd4* mutants

Tab 1: E13.5 *Adnp* cKO DEGs intersected with SFARI risk gene lists
Tab 2: E13.5 *Adnp* cKO DEGs intersected with SysNDD (definitive) risk gene lists
Tab 3: E13.5 *Adnp* cKO DEGs intersected with SFARI and SysNDD (definitive) risk gene lists
Tab 4: E13.5 *Chd4* cKO DEGs intersected with SFARI risk gene lists
Tab 5: E13.5 *Chd4* cKO DEGs intersected with SysNDD (definitive) risk gene lists
Tab 6: E13.5 *Chd4* cKO DEGs intersected with SFARI and SysNDD (definitive) risk gene lists
Tab 7: Gene list intersections: E13.5 *Adnp* cKO DEGs, E16.5 *Chd4* cKO DEGs, SFARI, SysNDD (definitive)

Original chemiluminescent scans of westerns shown in Fig. 1

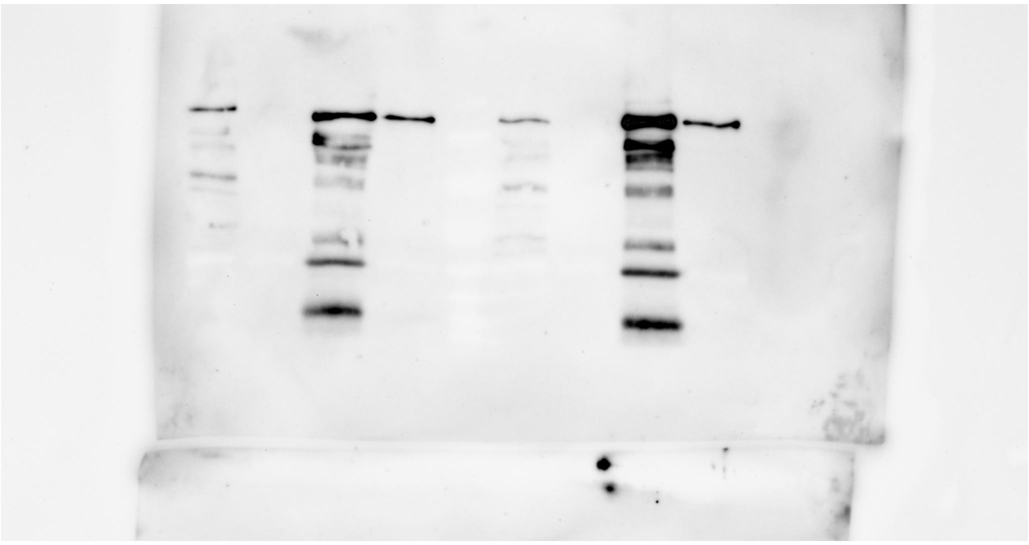

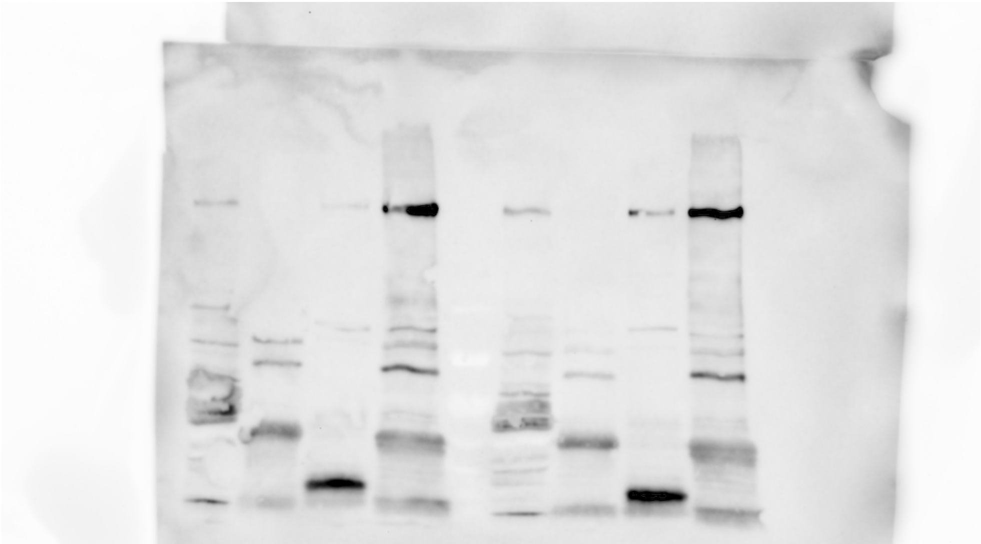

Original scans of Southern blots from Fig. S2

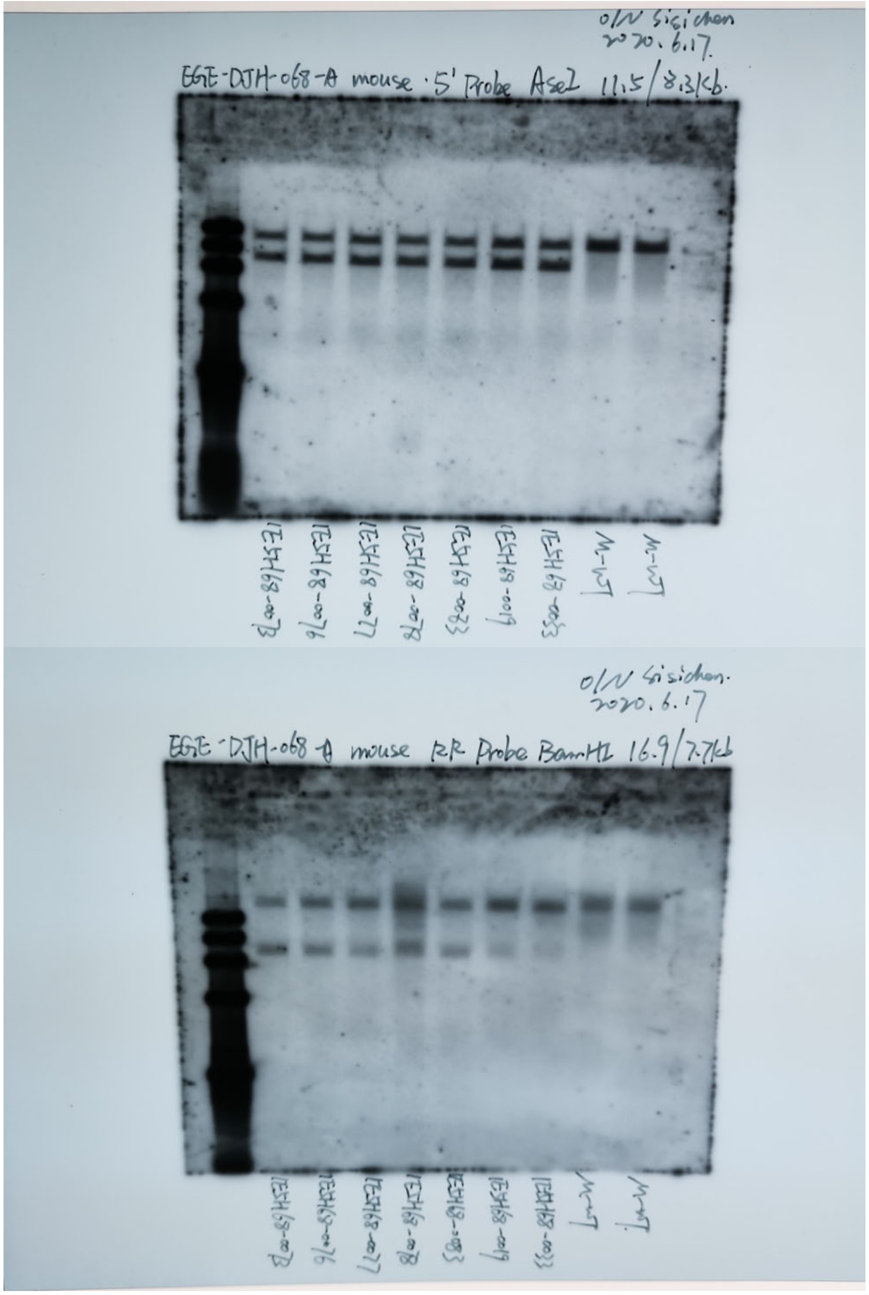
